# C.La.P.: Enhancing transformer-based genomic signal modeling by integrating DNA sequences and chromatin accessibility data

**DOI:** 10.1101/2025.02.19.638643

**Authors:** Panos Firbas Nisantzis, Carolina Gonçalves, Gonzalo de Polavieja

## Abstract

Transformers have shown promise in chromatin modeling but have primarily relied solely on reference DNA sequences, limiting their utility across multiple biological contexts. In this work we enhance transformer models by integrating reference sequences with ATAC-seq, a chromatin accessibility assay. Our Chromatin LAnguage Processing (CLaP) model combines a convolutional tokenizer, a transformer encoder, and task-specific components to predict multiple genomic signals from a single input. After pre-training with masked nucleotide prediction, CLaP achieved per-token F1 scores exceeding 0.8 for three target ChIP-seq assays in fine-tuning. Attention mechanism analysis revealed that CLaP detects CTCF binding sites with nucleotide-level precision by learning the sequence preference of the CTCF factor and the characteristic ATAC-seq patterns that are caused by protein binding events. Additionally, CLaP predicts protein-DNA binding events not captured by the ChIP-seq ground truth. These findings highlight CLaP’s potential to expand chromatin modeling by incorporating molecular assay data alongside sequence information.

## Introduction

### Cis-Regulation

Cis-regulatory elements (CREs) are non-coding regions of the genome that regulate the transcription of target genes. This is achieved through short sequence patterns that are contained in CREs, where Transcription Factors (TFs) preferentially bind. TFs in turn facilitate or inhibit the “Transcriptional Machinery” [1] from binding at the transcription start site of the gene and starting the process of transcribing DNA into RNA. CREs are highly dynamic. Their activity varies across different tissues or cell lines [2] and they are themselves controlled by complex regulatory pathways [3]. We are increasingly realizing the importance of CREs. In evolution they are a major driving force of diversity [4]. In development their precise dynamics in space and time are crucial for the healthy development of zygotes into multicellular organisms [5]. In health, CRE mutations can alter gene expression and contribute to human disease [6]. These regions contain chromatin alterations that differentiate them from the rest of the genome [7]. Molecular assays, followed by sequencing, are commonly used tools to detect and characterize CREs, with ChiP-seq [8] and ATAC-seq [9] being two popular examples (Fig 1A). ChiP-seq provides a genome-wide signal of protein occupancy, which can be used for detecting altered histones or TFs, while ATAC-seq offers a genome-wide signal of chromatin accessibility. While these tools are the current gold standard and invaluable in genomic research, and despite advancements in Next-Generation Sequencing techniques, molecular assays are still costly in terms of both time and materials. Furthermore, they often have limitations such as antibody availability [10] or TF binding site resolution [11].

**Figure 1:**
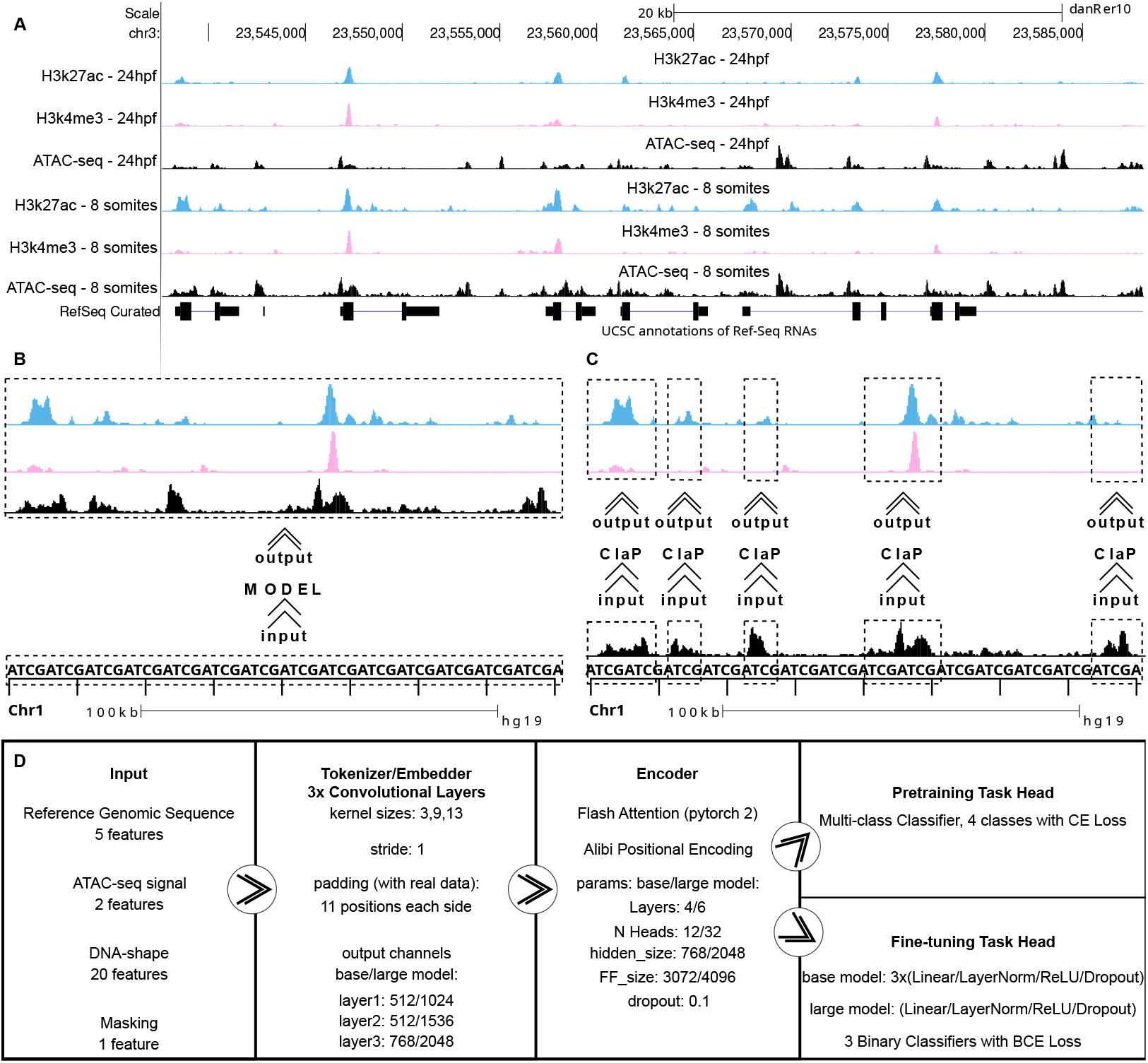
CLaP’s core ideas and architecture. **A**, Example screenshot of the UCSC genomic browser, typically used to browse genome-wide signals of cis-regulation. The figure shows the genomic signal of six genomic assays from two developmental stages of zebrafish (Danio Rerio), 8 somites and 24 hours post fertilization. In each stage, we see one ATAC-seq assay and two ChIP-seq assays of chromatin modifications (H3k27ac and H3k4me3). The browser is focused on a number of hox genes, marked in the bottom of the figure with black bars and lines. The peaks of the ATAC-seq experiments are the genomic regions that will become samples of our model, and the peaks of the histone assays is what the fine-tuned model will be trying to predict. **B**, The typical approach to modeling cis-regulation involves incorporating large ranges of the reference genome into a model that simulates all the contexts in which it has trained with. **C**, ATAC-seq is used to isolate Cis-Regulatory Elements. CLaP then incorporates ATAC-seq signal into its input. The model outputs a context-specific simulation of the genome-wide signals it was trained with. **D**, A simple diagram of the model’s overall architecture and overview of the hidden sizes used for the base and large versions of the model.

Research in the field of cis-regulation almost invariably relies on contrasting different biological contexts such as cell types, developmental stages, or tissues. As such, a great number of assays often need to be performed in order to gain a comprehensive understanding of the biological processes involved. Computational models can therefore be very useful in tasks such as getting initial results or selecting putative targets. In some cases, computational approaches are the only way to produce answers [12]. For example, the most popular molecular technique that we use to find where the CTCF protein is bound on the genome, a ChIP-seq against CTCF, produces statistically significant peaks of about 300 base pairs (bp) wide as a consequence of the molecular assay which aims to shear the chromatin to a target size of 100–300 bp [13]. The protein itself is up to about 30bp wide [14], so we are left with some uncertainty with regards to the exact binding site and for now this problem can only be approached with modeling methods.

### Transformers

Genomic sequences can be considered as a type of language and as such they have been featured in transformer research since its early stages [15]. Following their success in other fields such as protein folding [16], transformer models are now increasingly being used cis-regulation [17, 18, 19, 20].

Most transformer applications in cis-regulation are designed to generate multiple chromatin profiles using only the reference genomic sequence as input [21, 22, 23, 24, 25]. However, since the reference genomic sequence is immutable and shared by all cell types of any given organism, this design choice hinders a model’s ability to produce context-aware results. In BigBird [15], the model learns 919 binary classifiers of chromatin profiles, including 690 TF binding profiles for 160 different TFs, 125 DNase I sensitivity profiles and 104 histone-mark profiles. DeepMind’s enformer model [18] similarly relies on genomic sequence and outputs 5313 human and 1643 mouse profiles. BigBird’s output includes 100 different profiles pertaining to the CTCF TF, modeling 100 different experiments from cell lines or tissues. These models are restricted to the specific experimental contexts represented in the training data and are therefore unable to generalize to new, unseen conditions. A potential strategy to circumvent this limitation involves combining model predictions with data derived from molecular assays. For example, one could produce model predictions only for interesting genomic regions that have been identified through a molecular assay. Those predictions would be somewhat reliant on context, but if a molecular assay is to be performed, why not integrate it into the model itself? In a very recent work [26] ATAC-seq is used with a transformer, but only a single accessibility score value is kept per cis-regulatory region, thus not taking advantage of the per-nucleotide signal richness of the assay.

Another common choice in the literature is to design models that take as input the largest possible genomic region. The motivation for this is to allow the model to capture as much genomic information as possible, including distal CREs [18]. This is a reasonable motivation, but even impressive engineering only manages to fit about 200.000 bps as input for the model, when distal CREs are known to exist up to Mbps away from their target gene [27]. Furthermore, this approach includes in the input genomic sequences that are neither genes nor CREs. These regions might include repetitive sequences and transposon remnants, are typically not conserved and often have no known functional role [28] making the benefits of including them in the input doubtful.

One final drawback of this approach is that processing these very large input sequences is computationally prohibitive with the currently available hardware. As a result, inputs must be compressed to fit within memory limits, typically using max-pooling, which reduces resolution. This trade-off might be particularly important when attempting to derive biological insights with nucleotide resolution.

### A model of CREs

We present C.La.P. (Chromatin Language Processing), a transformer encoder model of cis-regulatory elements that overcomes the previously mentioned challenges. Our approach revolves around two core design principles: using training samples that are based on individual CREs (as predicted by ATAC-seq) instead of arbitrary long genomic spans, and integrating the signal of ATAC-seq in the model’s input (Fig 1B,C). ATAC-seq provides a genome-wide chromatin accessibility signal which is typically used to define putative CREs by identifying regions with significantly elevated ATAC-seq signal. We do this to define samples, but also integrate the per-nucleotide signal in the input of the model. Protein-DNA binding events leave ‘shadows’ on this signal, an observation that has long been used to model the binding of TFs on DNA [29, 30]. We expect that this integration will allow CLaP to model TF binding events in a context-sensitive manner, ultimately allowing it to comprehensively model the input CREs.

We pretrain and fine-tune this model with data from the ENCODE project and show that the finetuned model accurately predicts histone modifications and captures their dynamics in a very challenging multi-context dataset. Furthermore, the model identifies CTCF binding sites with nucleotide precision by de novo, discovering both the protein’s consensus motif as well as footprints on the ATAC-seq signal.

## Results

### Training

After 30 epochs, the baseline model achieves a per-token F1 score of 0.79 in the pre-training validation dataset. In the same amount of computing time, the large model only trains for five epochs, but achieves an F1 score of 0.82, having climbed above 0.8 and surpassing the baseline model after only two epochs (Fig. 2A). In the fine-tuning task (Fig. 2B), the large model reaches per-token F1 scores of 0.79, 0.84, and 0.8912 for CTCF, H3K27ac, and H3K4me3, respectively. Pre-training for more epochs leads to better fine-tuned models (Fig. 2C), but with diminishing returns. This information can help make informed decisions about the optimal pre-training duration, particularly when minimizing computational costs or time is a priority. The model mostly classifies uninterrupted segments of tokens as belonging to any of the three classes. Since, in the field of cis-regulation research, range-based analyses are more relevant than nucleotide-based ones, we also computed per-CRE F1 scores. With this we examine if CLaP manages to successfully classify ranges instead of individual tokens, across all three classes. In Fig. 3B (fine-tuning validation dataset), we apply this analysis per biosample and show individual experiments registering F1 scores of over 0.9 for all classes, while the mean F1 score between assays remains above 0.8 for all three classes.

**Figure 2:**
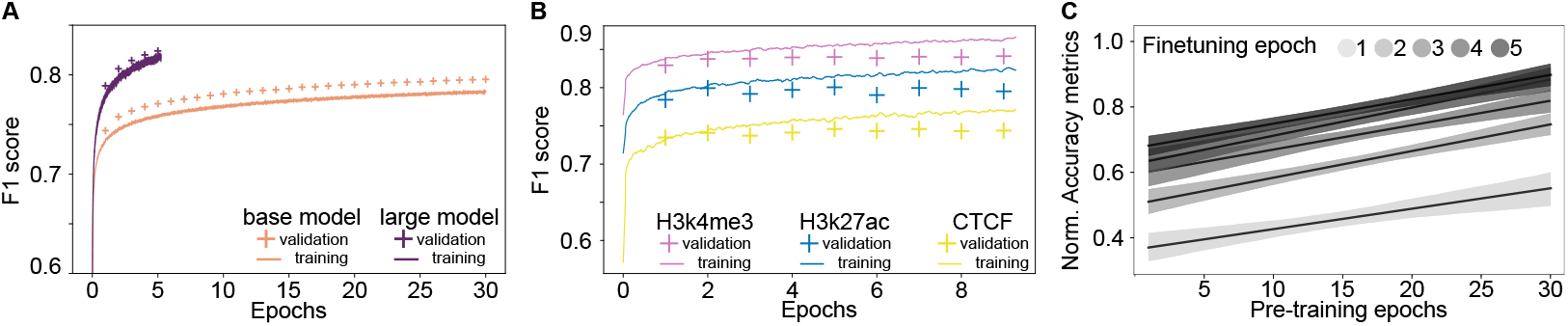
Pre-training and Fine-tuning. **A**, The pre-training progress of the base and large models. The continuous line tracks the F1 score (y axis) of the batches seen in the last 20 seconds of computation (note: dropout is on here). At the end of each epoch (one full pass over all training data) the model is evaluated on the validation subset, and the resulting F1 score is marked with a cross (dropout off). **B**, The fine-tuning progress of the large model. As in panel A, the continuous line tracks the metric during training time while the crosses mark the result of evaluating the model on the validation set at the end of each training epoch. **C**, Effect of pre-training depth on fine-tuning performance. The base model’s pre-trained weights at different epochs (x-axis) were used to fine-tune it for 5 epochs. Normalized accuracy metrics for the three fine-tuning classes were computed on the validation dataset (y-axis) and averaged across classes. Regression lines were fitted to this data and visualized here, to illustrate the relationship between pre-training depth and fine-tuning performance.

**Figure 3:**
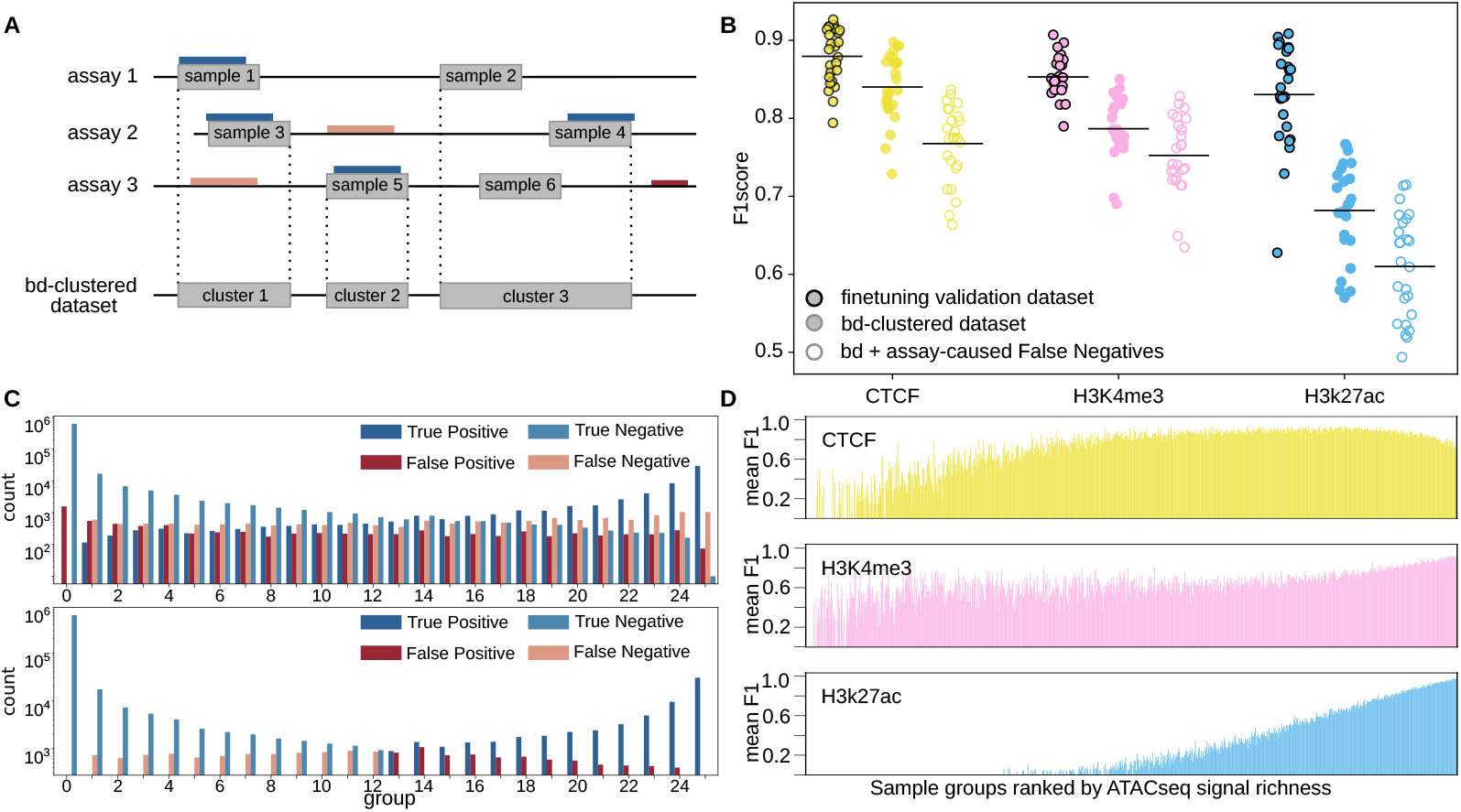
CLaP captures chromatin dynamics. **A**, The fine-tuning dataset consists of ATAC-seq peaks (gray bars) from a number of assays. Blue, pink, and red bars represent regions marked by ground truth ChIP-seq assays. Our samples might (in the case of blue bars), or might not (pink and red bars), overlap with ground truth regions from our target assays. The bd-clustered dataset (cluster 1-3) marks any region that was an ATAC-seq peak at least once in the dataset. Some ground truth regions (red bars) lie in regions that were not determined to be an ATAC-seq peak in any of the experiments of the dataset. **B**, F1 scores computed per class (CTCF, H3K3me3, H3K27ac), for each of the created datasets (fine-tuning validation dataset, bd-clustered dataset, bd + assay-caused False Negatives). The plot shows the F1 scores per biosample (individual marker), along with their respective mean (horizontal line). The ‘fine-tuning validation dataset’ is based on samples, while the bd-clustered dataset is based on clusters of samples that are evaluated in every assay. The third dataset (bd + assay-caused False Negatives) expands the bd-clustered dataset with a proportional amount of False Negative samples (see main text). **C**, Clusters from the bd-clustered dataset were grouped on the x-axis by the number of tissues (0–25) in which they are active for CTCF. The y-axis (log scale) shows TP, FP, FN, and TN predictions per group for CLaP and a hypothetical sequence-only classifier, as detailed in the main text. **D**, Average F1 score per group of samples ordered by increasing ATAC-seq signal richness. Samples were split into equally sized groups based on their mean ATAC-seq signal. Model performance declines in low-signal samples (left side of each subplot), particularly for H3K27ac, and shows a slight decrease in CTCF class for the richest-signal samples.

### CLaP captures chromatin dynamics

Our fine-tuning dataset is compiled with data from 25 different biosamples. ATAC-seq assays on those biosamples are used to define samples (see samples in Fig. 3A) while ChIP-seq assays on the same biosamples are used to define the target classes. Ground truth regions might (in the case of blue bars), or might not (pink and red bars), overlap with ATAC-seq peaks in their respective biosample. The cases where a ground truth region does not overlap with an ATAC-seq peak could be considered a False Negative (FN) when evaluating how well ATAC-seq and CLaP approximate the ground truth. To give our model a chance to recover these missed predictions, we compiled the bd-clustered dataset (Fig. 3A) by grouping our samples into overlapping regions with ‘bedtools cluster’ [31]. This produces a set of regions that have been marked with ATAC-seq in at least one assay. We evaluated the model on these regions in every available biosample and plotted the mean performance per biosample in Fig. 3B (bd-clustered dataset).

Interestingly, there are still ground truth regions that do not overlap with this dataset (Fig. 3A, red bar). This is caused by binding sites of CTCF that, for some reason, are never registered as ‘open’ in any of the ATAC-seq experiments. We included a proportional amount of these regions (10% of total, matching the ratio of validation data to the full dataset) as FN to the bd-clustered dataset to produce the third and harshest evaluation of Fig. 3B (bd + assay-caused FN). Even under these very strict conditions, the model achieves F1 scores of over 0.8 in many experiments of CTCF and H3k4me3 and the mean F1 score for those two classes is over 0.75.

Regions or clusters in the bd-clustered dataset (Fig. 3A) may overlap a ground truth region in zero, one, or multiple contexts. Rarely activated, tissue-specific regions are active in a few contexts while ubiquitous, always-on regions are active in all contexts. In Fig. 3C (see Supp.Fig. 2-3 for the H3K27ac and the H3K4me3 classes), we split these regions into groups, based on how many contexts they are active in. If a region never overlaps with a target it’s included in group 0, if it overlaps with a target in all 25 contexts it is put in group 25, and so on. We then computed and plotted the per-CRE True Positives (TP), False Positives (FP), True Negatives (TN) and FN counts of the model for each group.

As a reference model, we use a theoretical sequence-only model that would achieve the best possible accuracy in this dataset. A sequence-only model cannot make differential predictions between the various contexts of this dataset and consequently would classify each sample in exactly the same way between contexts. Thus this hypothetical model would achieve its best possible accuracy, if it classifies regions as “always active” when they are active in at least 13 contexts and “always inactive” when they are active in less than 13. Any other choice would produce more wrong than correct predictions and would still be naively classifying all regions as always-active or always-inactive. This hypothetical model can nevertheless achieve high accuracy or F1 scores, but fails to make any TP predictions for half of the dataset, where the interesting tissue-specific CREs are found. Similarly, CREs that are open in the majority of contexts but are deactivated in a minority will be ignored by this model. It would essentially classify each genomic region as being always on or always off, completely missing any differential dynamics. In contrast, CLaP produces a high number of TP and TN predictions across the entire dataset, effectively capturing differential dynamics even in challenging multi-context analyses.

### CLaP integrates both sequence and ATAC-seq

#### ATAC-seq signal improves model performance

The ATAC-seq signal appears to be integral to the function of the model already from the pre-training stage. When we attempted to pre-train a version of the model while withholding the two ATAC-seq features (See Supp.Fig 1), the model failed to achieve F1 scores significantly higher than 0.5.

To further investigate the effect of the ATAC-seq signal in the model’s performance, we ranked the samples based on their mean ATAC-seq signal and split them into equally sized buckets (Fig. 3D). When the ATAC-seq signal is low or sparse (left side of the x-axis), the model relies more heavily on the genomic sequence for its predictions, leading to a decline in performance. The effect is most noticeable in the H3k27ac class, possibly due to the fact that the histone modification that is measured by the H3K27ac assay and the chromatin accessibility that is measured by ATAC-seq are often not synchronized; the chromatin of a region first gets modified and only becomes accessible later. Interestingly, in the case of the CTCF class, the regions with the highest amounts of ATAC-seq signal also suffer slightly in performance. A possible explanation could be that the samples with very high ATAC-seq coverage contain artifacts of the sequencing process that survived the data pre-processing and cleaning pipelines (low quality reads, duplicates, among others) all of which make the input noisy and confuse the model particularly in the case of CTCF, where the model is ‘looking for’ a single protein binding spot instead of broader chromatin patterns.

#### CLAP’s attention heads show language-like patterns

Next, we analyzed the model’s attention mechanism. This is a core component of the transformer architecture and through its analysis we tried to identify the specific elements of the input data that the model prioritizes when making predictions. Transformer models have a number of layers, each containing a number of independently trained heads, with each head learning a different attention matrix.

The attention matrix has as many rows and columns as the number of tokens of the input tensor and maps the relationship of each token against all other tokens. The attention matrix varies from head to head and from one encoder layer to another, with each head potentially learning different patterns or aspects of the input sequence. In language models, these patterns can correspond to various linguistic features, such as syntax, semantics, or coreference [32].

Visualizing the self-attention matrices of individual heads from different layers of CLaP produces patterns similar to those observed in language models (Fig. 4 D). Diagonal patterns are the most common, but strong vertical lines, blocks, and heterogeneous patterns are also observed.

**Figure 4:**
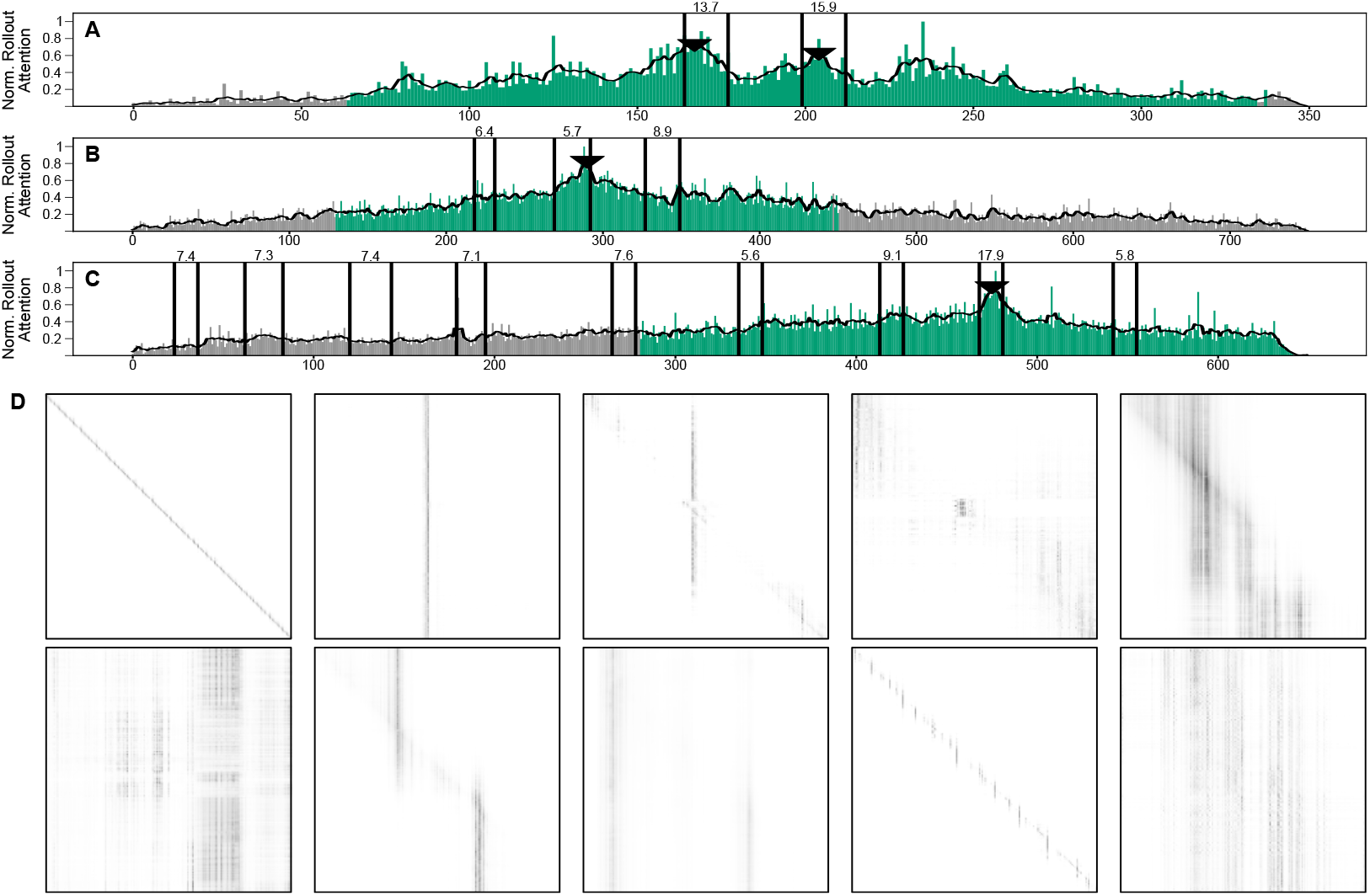
Investigating CLaP’s attention. **A,B,C**, Normalized attention rollout (y-axis) for three distinct samples (x axis is the token position on the sample). Bars show per-token attention values, which were smoothened by a moving average filter (black line). Green bars mark tokens classified as CTCF-positive, vertical black lines indicate CTCF PWM hits (scores above), and black triangles denote ‘mA attention’ peaks. **D**, Examples of self attention matrices from hand picked heads. These matrices measure how much attention is paid from each token to each other token. Diagonal patterns where tokens mostly pay attention to their own self and nearby neighbors are the most common. Vertical lines and focal points that remind patterns observed in language models often localize over CTCF PWM hits.

In particular, a number of heads (the third,fourth,and fifth in the case of the base model) show very strong patterns that colocalize with CTCF Position Weight Matrix (PWM) hits (a.k.a. ‘CTCF motifs’) (see Supp.Fig.5 and the Supplemental CTCF examples pdf file for some ‘raw’ examples). PWMs are short, sequence-only, kernel models that capture the protein’s sequence preference. They are the most commonly used model of TF binding. Clearly then suggesting that CLaP has learned this sequence preference de-novo.

We then identified CTCF PWM hits inside ground truth positive CTCF regions and stacked attention matrices centered on these hits (Sub. Fig. 6). To reduce noise in this signal, we subtracted from it stacked attention heads from random genomic positions. This amplified each head’s reaction to real binding events and revealed a number of striking patterns of activation. Are these activations just tracking the genomic sequence or do they also take into account the ATAC-seq signal? To investigate, we compared the activations of the individual heads in pairs of samples with identical sequences but different ATAC-seq signals (Supp.Fig. 4), i.e. we took the same genomic region from two different ATAC-seq experiments and evaluated them with the model. The self-attention heads whose activation relies heavily on the genomic sequence will produce highly correlated values since the genomic sequence is identical, while heads that rely more on the ATAC-seq will have lower correlations.

We saw both of these cases in our analysis (Supp.Fig.5) and crucially noticed that some heads (e.g. layer3|head3) produce strong activation patterns that colocalize with CTCF PWM hits while having relatively low skew score, meaning that their activation depends to a decent degree on the information of the ATAC-seq signal.

#### Attention rollout

To combine all of the model’s attention heads for a single sample, we employed the attention rollout method [33], which aggregates the attention matrices throughout all the layers and heads of the encoder. This approach generates an array of attention values spanning the entire sample as a measure of the overall attention allocated by the model to each position (or token) within the input sequence.

The resulting arrays often contain discernible peaks (Fig. 4 A-C), which we call maxAttention positions (mA positions). In the samples where the baseline model makes a positive prediction for CTCF, 98% of mA positions fell inside CTCF PWM hits. Performing the same analysis, i.e. naively taking the position of maximum attention inside ranges where the model is predicting CTCF (the highest green bar in Fig.4 A-C), we found a lower percentage of overlap with CTCF PWMs (around 60%). Nevertheless, the large model’s attention continues to peak over CTCF motifs. By applying a smoothing function to the model’s attention-rollout array across the sample (black line in Fig 4 A-C) and performing peak-calling on the smoothed array, we can identify more refined mA positions (black triangles in Fig 4 A-C) and increase their precision. With this method, the overlap between the large model’s ‘mA positions’ with CTCF motifs increases to 85%.

Although the model demonstrates improved precision and the capacity to detect multiple mA positions within a single sample (Fig. 4A), several PWM hits remain uncovered by attention peaks (as illustrated in Fig. 4 A-C). Typically, the model generates a single prediction that aligns with the highest scoring PWM hit (Fig. 4 C), but this is also not consistently observed across all cases (Fig. 4B). However, we do not interpret this as a limitation of the model. On the contrary, many PWM hits lack reliability, and an effective model should be capable of disregarding these while prioritizing the true positive instances. These results indicate once again that the model is capturing more than just genomic sequence patterns.

### CLaP detects CTCF with nucleotide resolution

For the CTCF class, the majority of per-token FP and FN predictions of the model happen at the edge of the target region. To demonstrate this, we selected samples that contained a ground truth or predicted CTCF, and aligned them on their centers (Fig. 5A, black arrows). The ground truth regions are derived from CTCF ChIP-seq assays. The actual target of these molecular assays, the CTCF proteins, are about 30 base-pairs long [14], yet the ground truth regions that we derive from the sequencing assays (our ground truths) are much bigger. In the case of ChIP-seq for CTCF, the suggested target size for DNA fragments is 200-250 nucleotides long [34] and as a consequence the derived genomic ground truth regions will have an expected mean width of more than 200bp. This is an unavoidable consequence of the ChIP-seq pipeline, the real target lies somewhere (usually near the center) inside the ground truth sequence, but its signal is defused in a noisy way around it. The very low FP rate at the center of the targets (Fig.5A orange line) suggests that the model is capable of detecting the actual CTCF binding sites, but has difficulty in correctly predicting how far the signal extends. This difficulty is expected, since this extension of the signal is random and the model has no means to correctly predict it. The sharp enrichment of CTCF ‘motifs’ or PWM hits in the center of this visualization also lends support to the argument that the model is detecting binding sites in order to accomplish the classification task. One more clue in support of this argument, comes from the fact that CTCF classification scores are the worst of the three classes when computed per-token, but become the best of the three classes when computed per-CRE. The model is correctly predicting binding sites, but loses accuracy when trying to predict the noisy signal extension around those sites.

**Figure 5:**
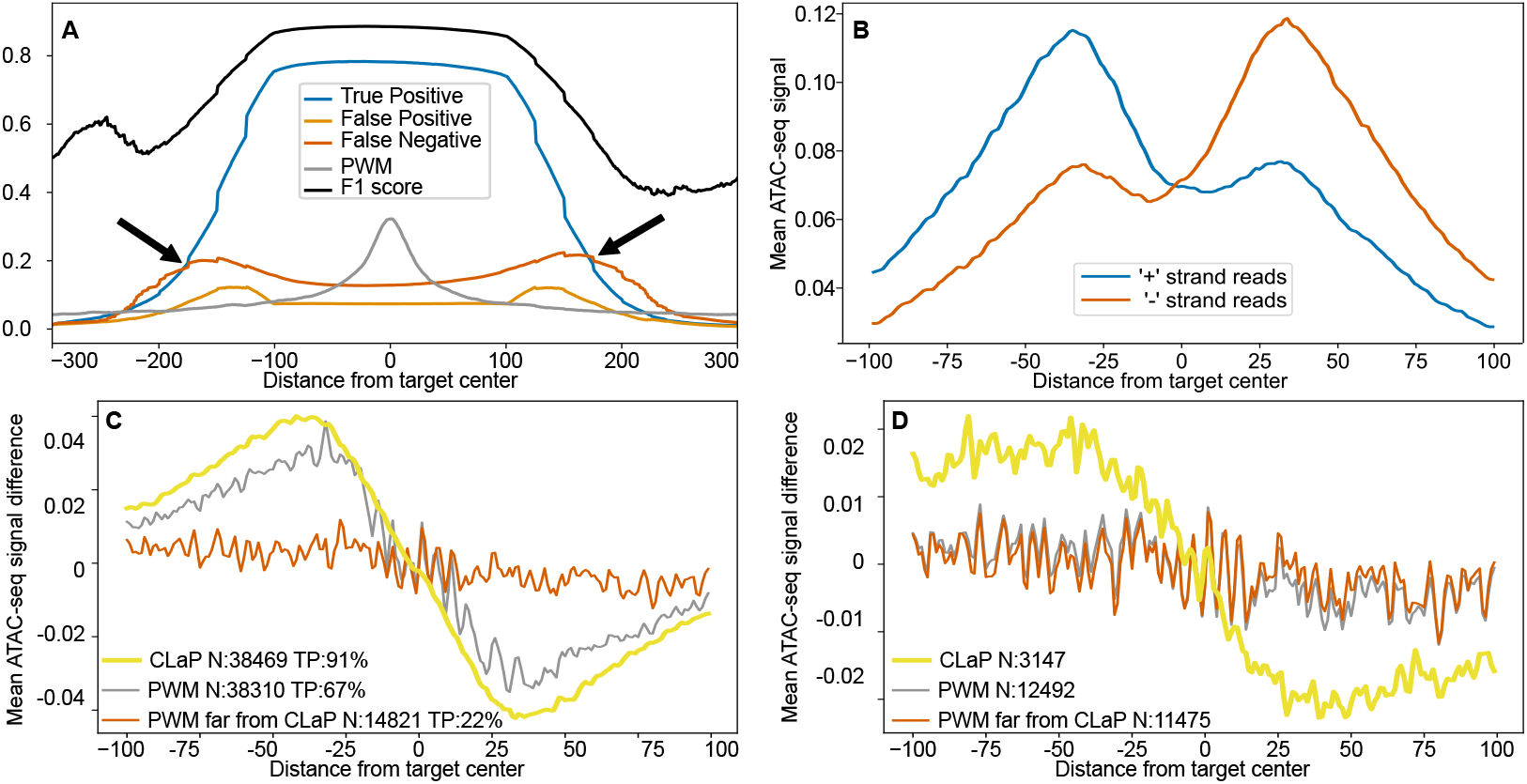
CLaP detects CTCF binding events. **A**, Samples with tokens that were classified as positive for CTCF either in the ground truth or the model’s predictions were aligned on their center and, for each position (here on the x axis), TP, FP,FN and F1 scores were calculated. The ratio of tokens overlapping CTCF PWM hits is also plotted as PWM. Black arrows point to ‘bumps’ in the FP and FN signals at a distance from the center. Samples with tokens classified as CTCF-positive (ground truth or model predictions) were center-aligned and TP, FP, FN, F1 scores, as well as PWM overlap ratios were calculated across token positions (x-axis). Black arrows highlight FP and FN “bumps” at a certain distance from the center. **B**, CLP peaks were collected for the entire validation dataset and aligned on their centers. For each position around the center (here on the x-axis), we plot the mean value of ATAC-seq signal, keeping the signals of both + and - strands separate to showcase the characteristic imbalance that is observed around protein-binding events in ATAC-seq experiments. **C**, Plotting the normalized difference between “+” and “-” ATAC-seq signals allows us to compare sets of positions. Here we compare CLP peaks and PWM hits with score higher than 11.5, both found inside the samples of our validation set. The third set, PWMs far from CLAP, is a subset of the PWM set containing only the PWMs that did not overlap a CLP peak. In each case, True Positives are the predictions that overlap with peaks from our ground truth CTCF ChIP-seq data. The predictions of CLaP maintain a slightly stronger ASI, while having a much higher TP rate. **D**, Like in panel C, but this time we compare predictions that would be considered False Positives based on overlap with the ground truth CTCF ChIP-seq. CLaP predictions still show some ASI here, suggesting that the model captures real binding events that were missed by the ground truth assay.

#### Protein occupancy through ATAC-seq

As we saw, the rollout attention of the model peaks over CTCF PWMs when it predicts a CTCF site. A more straight-forward and computationally effective way of extracting predicted sites from the model is to take the probability logits for the CTCF class, before softmax, to smoothen that signal and then look for peaks with scipy’s find_peaks function. These predicted sites, which we’ll call CLP peaks, also tend to overlap with CTCF binding sites, with half of them being less than 6bps away from the center of the nearest PWM (Supp.Fig.6). We analysed CLP peaks for known and de-novo PWMs (Supp.Fig.7). Among the known ones, two motifs of CTCF were the most highly enriched. From the de-novo PWMs, the top two were similar to the known CTCF PWMs. By plotting the signal of ATAC-seq around CLP peaks, and keeping the “+ mapped” and “-mapped” sequencing reads separated, we observe evidence of protein occupancy (Fig. 5B). There is a clear depletion of signal at the center, where the protein is expected to be bound and thus protecting DNA from cleavage. Furthermore, the + and - reads display the characteristic imbalance that is observed around protein binding sites [30]. This is solid, molecular-assay-based evidence that the putative binding sites are real ones. It is not easy to quantify, especially for individual sites, but it offers a way to compare groups of putative binding sites.

By subtracting the “- strand” signal from the “+ strand” signal, we can visualize the imbalance in a single line and make comparisons. The steeper the decline around 0, the higher the protein binding for a given set of genomic sites. In Fig.5 C, we compare CLP peaks to PWM hits in our validation dataset. We set a relatively conservative threshold of 11.5 for the score of PWM hits in order for this method to produce a comparative amount of predictions to those of CLaP. This is the kind of predictive performance you would expect by applying these two methods in a dataset of ATAC-seq peaks (putative CREs) from multiple different contexts. CLaP’s predictions show much higher TP rates and stronger protein binding signal. Removing the PWM hits that coincide with CLP peaks (‘PWM far from CLaP’) shows how poor the protein binding signal is over sites that CLaP does not select.

In Supp.Fig. 8, we compare only the TP predictions from CLaP and from PWMs. These are predicted binding sites that have been validated through the ChIP-seq assay, but through the ATAC-seq imbalance approach we can evaluate them further. In this scenario, CLaP retains the same amount of ATAC-seq imbalance for a significantly larger amount of predictions. Additionally, the subset of PWM hits that do not overlap with CLaP predictions again show a significantly lower amount of ASI.

Finally in Fig. 5D, we looked at predictions that fell outside the GT, i.e. the “FPs”. In comparison to the PWM FP sites, which are many more and display almost non-existent ASI, our model’s predictions display some ASI indicating that a number of these predictions are real binding events that were not captured by the ChIP-seq assay.

## Discussion

We developed a model of cis-regulatory elements (CREs) that integrates prior biological knowledge with modern machine learning (ML), leveraging recent advances in transformer architecture. Transformers have shown strong performance in sequence modeling, making them well-suited for genomic applications. We reasoned that incorporating chromatin accessibility signal (ATAC-seq) alongside the reference genomic sequence would enhance CRE modeling.

A key challenge in genomic modeling is the limited availability of comprehensive biological assays. While paired datasets containing ATAC-seq and other assays are scarce, ATAC-seq alone is more widely available. To address this, we implemented a pretraining and fine-tuning scheme: the model is pretrained using ATAC-seq data in a semi-supervised manner to learn a foundational representation of chromatin accessibility before fine-tuning on smaller, task-specific datasets. This approach allows the model to generalize well despite the scarcity of fully labeled datasets. Our results indicate that models trained solely on fine-tuning tasks, without pretraining, perform poorly—barely exceeding baseline performance. In contrast, pre-training significantly improves downstream accuracy, with longer pretraining yielding better results, albeit with diminishing returns beyond a certain threshold. These findings highlight the importance of leveraging abundant accessibility data for pre-training in genomic ML models. In addition to using ATAC-seq as an input to CLaP, we leveraged it to define individual samples. Each sample corresponds to a genomic region identified as ‘open’ through the ATAC-seq pipeline, allowing the model to focus on the most relevant cis-regulatory regions in each biosample.

For fine-tuning, we trained CLaP to predict genomic signals from three distinct ChIP-seq assays: CTCF, H3K27ac, and H3K4me3. These assays capture different aspects of chromatin state—CTCF binding sites define insulators and regulatory boundaries, H3K27ac marks active enhancers and promoters, and H3K4me3 is associated with transcriptionally active promoters. By learning these signals, the model gains deeper information about chromatin organization and gene regulation.

One of the aims of CLaP is to replace expensive ChIP-seq assays with ATAC-seq-based inference. If the model performs optimally, users would only need to perform the relatively inexpensive ATAC-seq assay, and CLaP could accurately infer the chromatin state across multiple regulatory marks. In our validation dataset, spanning 25 biosamples from 16 tissues, CLaP achieved mean F1 scores of 0.88, 0.83, and 0.85 for predicting CTCF, H3K27ac, and H3K4me3 signals, respectively (Fig. 3B), demonstrating its robustness in nucleotide-level classification

We found that CLaP’s predictions were less accurate for tokens or regions marked with the H3K27ac histone modification. This limitation arises because not all genomic signals co-occur precisely with ATAC-seq peaks. H3K27ac is associated with both poised [35] and active enhancers. While active enhancers exhibit strong ATAC-seq signals due to open chromatin, poised enhancers are primed with histone modifications but remain largely inaccessible, resulting in weak or absent ATAC-seq signals. As a result, CLaP is forced to rely more on genomic sequence alone in these regions, making it susceptible to the known limitations of sequence-based models, such as reduced ability to distinguish between functionally distinct but sequence-similar elements.

Despite this limitation, CLaP successfully captures the activation and inactivation of cis-regulatory elements across a multi-tissue dataset. It also identifies rare enhancer activations and inactivations (Fig. 3C) that sequence-only models cannot detect, highlighting the importance of integrating contextual biological information.

CTCF binding differs from histone modifications and offers a useful test case for the model’s ability to capture TF binding events, which lies at the core of modeling the mechanics of cis-regulation. While histone modifications cover broad genomic regions, CTCF ChIP-seq marks binding events of individual proteins. CTCF binds DNA with sequence specificity, typically modeled with PWMs. Though ChIP-seq-derived target ranges span 200–400 bp, the actual binding site is only 30 bp. When bound, CTCF protects DNA from cleavage, creating a footprint in the ATAC-seq signal. This footprint, reflected as a local depletion of accessibility within a broader accessible region, serves as an important feature for the model, allowing it to distinguish direct CTCF binding from mere sequence similarity.

The model’s raw probability scores, aggregated attention, and specific attention heads create peaks that strongly overlap with CTCF PWMs and ATAC-seq signal depletion. This suggests that, even with the inherent noise in ChIP-seq-based CTCF annotations—arising from crosslinking biases and broad peak calls—the model effectively captures direct protein binding events. CLaP achieves this by integrating sequence preference with chromatin accessibility features. Rather than relying solely on sequence motifs, the model also leverages ATAC-seq footprints, which reflect regions protected from transposase cleavage upon protein binding. This dual mechanism enables CLaP to distinguish functional CTCF binding sites from sequence motifs that lack biochemical evidence of binding.

To extract CTCF binding predictions, we identified peaks in the model’s raw probability outputs (CLaP peaks). Examining the reference sequence around those positions, we found that both known and denovo PWMs for CTCF are enriched, adding to the evidence that the model captures sequence motifs associated with binding.

Additionally, ATAC-seq signals around CLaP peaks show characteristics of protein binding, including central depletion (Fig. 5B) and strand-specific read imbalance (Fig. 5C, D). Some of the model’s attention heads specialize in detecting these patterns across both the genomic sequence and ATAC-seq signal. Activation patterns of these heads are centered around known CTCF PWMs.

CLaP peaks had a higher true positive rate than PWM hits and exhibited stronger ATAC-seq signal imbalance (ASI) (Fig. 5B). ASI, a qualitative proxy for protein binding, allows comparisons of putative CTCF sites without relying on ChIP-seq data. PWM hits within ChIP-seq target regions but outside CLaP peaks showed significantly weaker ASI (Fig. 5C), while CLaP peaks outside ChIP-seq ground truth still exhibited non-zero ASI. This suggests that some CLaP predictions represent true binding events missed by ChIP-seq.

Performance could improve with a more comprehensive training dataset. Our current approach required biosamples with four molecular assays, as we trained on all three classes simultaneously. A model focused solely on CTCF could leverage a larger dataset. Additionally, increasing model size could yield further gains, as larger versions of CLaP significantly outperformed baseline models.

CLaP models individual CREs, but higher-order chromatin dynamics, such as chromatin architecture and transcriptional regulation, emerge from interactions among multiple CREs. Extending CLaP by integrating predictions across neighboring CREs into a meta-model could extend its abilities to inferring chromatin interactions, enhancer-promoter coupling, and broader regulatory landscapes. This approach could bridge local sequence-based predictions with larger-scale chromatin organization, providing a more complete view of gene regulation.

## Supporting information

Supplemental CTCF examples

## Glossary

ASI: ATAC-seq Signal Imbalance
ATAC-seq: Assay for Transposase-Accessible Chromatin using sequencing
BCE Loss: Binary Cross Entropy Loss
bp: Base Pair(s)
CE Loss: Cross Entropy Loss
ChIP-seq: Chromatin Immunoprecipitation followed by sequencing
CLaP: Chromatin Language Processing
CLP: CLaP-derived peak (based on raw probability output)
CRE: Cis-Regulatory Element
CTCF: CCCTC-binding factor
ENCODE: Encyclopedia of DNA Elements
F1 Score: A statistical measure of a test’s accuracy (harmonic mean of precision and recall)
FN: False Negative
FP: False Positive
H3K27ac: Histone 3 Lysine 27 Acetylation
H3K4me3: Histone 3 Lysine 4 Trimethylation
mA: MaxAttention (peaks of CLaP’s attention rollout)
Mbps: Megabase Pair(s)
ML: Machine Learning
PWM: Position Weight Matrix
TFs: Transcription Factors
TN: True Negative
TP: True Positive

## Online Methods

### Datasets

We obtained data, (.bam and .bed files), from the ENCODE project’s portal. For the pre-training task, 75 ATAC-seq experiments were selected (See Supplemental Table 1).

For the pre-training task, each ATAC-seq peak from each experiment became a sample. These peaks overlap on the reference genome, so in order to avoid testing the model on genomic positions that it had ‘seen’ in the training, we grouped overlapping peaks into groups, or ‘clusters’ with ‘bedtools cluster’ [31]. For the fine-tuning task, using the same training/validation/test split as in the pre-training dataset, we selected the subset of samples where, besides ATAC-seq, the available data included CTCF ChIP-seq, H3K27ac ChIP-seq, and H3K4me3 ChIP-seq as well (See Supplemental Table 1).

For ATAC-seq data, we download the raw .bam alignment files and use a de-biasing tool [36] to produce genome wide signal .bigWig files of plus-aligned and minus-aligned reads. We also download the less strict ATAC-seq peaks (IDR ranked .bed files), as produced by the ENCODE project to use for sample creation.

For ChIP-seq data, we download the stricter (IDR thresholded .bed files) peak set, as produced by the ENCODE project.

The ENCODE4_v1.9.1_GRCh38.fasta genome was used wherever needed.

### Input preparation

For pretraining, each ATAC-seq peak from each experiment was a sample. Samples from various tissues were merged in the same dataset here and the model was ignorant of the biological context from where each sample comes from.

For each sample, we collect the relevant reference genomic sequence (6 features: A,C,T,G,N,MASK), the relevant ATAC-seq signal (2 features: + and - mapped reads are kept separate since their imbalance carries contextual information), as well as sequence based DNA-shape features as produced in [37] (20 features). The total number of features was 28. Convolution layers shrink the input so to maintain the input sample at the specified length, a small amount of real-data padding is added to the input.

The DNA-shape features were normalized using sklearn.preprocessing.StandardScaler and further with MinMaxScaler to stay in the [0-1] range.

ATAC-seq genome-wide data were de-biased and their values are also normalized between 0 and 1. To normalize, we computed the 99.9 quantile highest single nucleotide ATAC-seq value in the training subset. Every other nucleotide’s value (in all dataset) is clipped to this maximum value, and divided by it in order to bring the range of ATAC-seq values between 0 and 1.

### Training tasks

While the combination of genomic sequence and ATAC-seq signal can have a wide and versatile application range, training a machine learning model to simulate other genomic signals as its output means you need datasets where ATAC-seq and the other, target, assay have both been performed on the same biosample, ideally from the same laboratory. This drastically reduces the amount of available datasets, leading to relatively fewer available training samples which in turn can lead to overfitting models, as the models do not have sufficient data to learn robust patterns and rules that generalize across different genomic sequences and assays.

However, a first lower-level understanding of the genomic patterns, the dynamics of ATAC-seq signals and the interplaying between these can form the basis for learning more specialized, higher-level features of the chromatin at a secondary stage. To gain this lower-level understanding a model can be trained without target assay data, on a dataset and task that are derived from the more readily available input data. This first stage of learning is called pre-training and allows us to build a broad versatile model, with a rich, generalizable understanding of the genomic landscape. We call this a foundational model. At a later stage the model can have a secondary round of training, this time on the smaller datasets that include target assays, in order to specialize in more important tasks.

Drawing inspiration from the BERT model [38], we pre-trained our model on a token (nucleotide) masking task. This allows us to train on unlabeled data and exploit the bidirectional context of genomic sequences. At this stage, we expect that the model can potentially discover inherent structure in signal behavior, along with underlying patterns in the genome, such as sequence motifs, transcription start sites, and chromatin accessibility.

We used the same masking procedure as in [38], although adding some supervision to the way the tokens to be masked are selected. First, to promote a more evenly distributed and representative selection of masked tokens, we divided the input sequence into 6-nucleotide buckets. For each bucket, we choose one nucleotide to be masked. The selection process is as random as possible, but follows some criteria to guarantee that potentially relevant genomic positions are preferentially selected:

1. Pick a position randomly from the set of nucleotides associated with Single Nucleotide Polymorphism (SNPs) [39].
2. If there are no nucleotides associated with SNPs in the bucket, choose one that is conserved or accelerated [40].
3. If no nucleotides are conserved or accelerated, just randomly choose one of the 6.
4. Additionally, if an absolute genomic position has been masked previously in the dataset, it is skipped and another one is chosen instead.

Although this results in a fixed set of masked tokens for the whole dataset, this selection process enhances the pre-training task by challenging the model to predict the more meaningful nucleotides. It also prevents the model from focusing too much on more uninformative regions of the genome and learning irrelevant features.

After selecting all tokens per sequence, the masking occurs by treating these tokens as ‘N’ nucleotides and giving them DNA-shape features accordingly. This way the model cannot use the shape features to deduce the masked nucleotide. Their ATAC-seq signals are also set to -1. The model is then tasked to predict these masked tokens’ nucleotides.

The pre-trained model is then used for fine-tuning on three binary classification problems, in a supervised learning setting. For each token of the non-padded input sequence, the model is tasked to predict if that genomic position is overlapping a ‘peak’ in ChIP-seq, H3K27ac ChIP-seq, and H3K4me3 ChIP-seq.

### Model architecture

C.La.P architecture can be divided into three main parts: the Tokenizer and Embedder; the Encoder, and the Task Head (Fig.1 D). The model takes as input a sequence with a maximum length of 1800 positions, where each of its positions corresponds to a single nucleotide of a genomic sequence. In contrast with other transformer-based models that tokenize the dataset sequences a-priori [15, 22], defining a fixed vocabulary of nucleotide sets, we decided to learn these instead. By training our own tokenizer, we aim to learn relevant and dynamic genomic patterns that may change with context. We leverage convolutional layers to exploit local genomic patterns in the full sequence, independently of their position, splitting it into tokens and representing these in high-dimensional vectors which we call embeddings.

Our Tokenizer and Embedder consists of 3 convolutional blocks. Each block has a 1D Convolutional Layer with no bias, followed by a Batch Normalization layer and an activation function (in this case, we used ELU [41]. These convolutional blocks have stride of 1 and kernel sizes of 3, 9 and 13, respectively. This results in a total receptive field of 23 positions. As our input was previously padded with 11 nucleotides on each side, an embedding vector is produced for every nucleotide of the original input sequence. Each embedding represents a window of 23 consecutive nucleotides centered on its respective nucleotide position. Our tokens are therefore dynamic 23-length overlapping windows of genomic patterns that aim to exploit relevant local contexts for every position in the genomic sequence.

C.La.P’s Encoder is a multi-layer bidirectional transformer encoder responsible for learning the long-term dependencies of the tokenized and embedded local patterns. Our implementation follows the original one (see [42] for the full architecture), except for the positional encoding and the addition of Flash Attention for faster performance [43]. We use a relative positional encoding called Alibi positional encoding [44] instead, which takes into account information about the relative distances between each pair of tokens when learning their global interactions.

The transformer processes the full embedded sequence and outputs a new high-dimensional representation for each of its positions. These are the input of the Task Head, which allows us to train the model for different tasks. Here, we set two tasks (pre-training and fine-tuning), both as a classification problem carried out by a final Linear Layer. The latter is preceded by at least one block of one Linear Layer, an activation function (GELU for pre-training and ReLU for fine-tuning). The further details of this architecture are described in Section Model Configuration.

### Training and evaluation

#### Model configurations

C.La.P has two configurations used in this study: the base model and the large model. Both configurations share the same strides (S) and kernel sizes (K) for the three convolutional layers, differing only in their hidden sizes, the transformer’s hyperparameters and Task Head’s architecture (Fig. 1D). The kernel sizes were specifically chosen to ensure a receptive field large enough to capture genomic patterns found on TF binding sites [45], covering 23 nucleotides (as outlined in the *Model Architecture* section above). Particular attention was also given to the first two kernels to ensure these were small enough to capture simpler, shorter patterns that contribute to these motifs [46]. The remaining model’s parameters were firstly selected within the ranges encountered in the literature [38] and then empirically tuned based on performance and memory constraints.

Here we denote the number of layers or blocks as L, the hidden size as H, the number of self-attention heads as A and the hidden size for the feed-forward block in the transformer as FF_H. We report the following model configurations: C.La.P base model (Tokenizer and Embedder: L=3, S=1, K=3,9,13, H=512, 512, 768, Dropout=0.1; Transformer Encoder: L=4, H=768, A=12, FF_H=3072, p=0.1) and C.La.P large model (Tokenizer and Embedder: L=3, S=1, K=3,9,13, H=1024, 1536, 2048, Dropout=0.1; Transformer Encoder: L=6, H=2048, A=32, FF_H=4096). The former represents the data in a 768-dimensional embedding space and the latter in a 2048-dimensional one.

Depending on the task and model configuration, we have a slightly different Task Head architecture. Here, we further denote the number of classes as N, which matches the output dimension of the last linear layer of each head. We report two head configurations per task: pre-training Head (L=1, H=768 or 2048, N=4) and fine-tuning Head (L=3 or 1, H=768 or 2048, N=3) for base and large models, respectively. We also added Layer Normalization to each layer of the latter task head to help stabilize training.

We end up with two model sizes per task. For the pre-training, we have a base model with a total number of 36M parameters and a large model with 596M parameters. For the fine-tuning, these model configurations have 37M and 596M parameters, respectively.

#### Training details

We split the pre-training dataset into 80% of the clusters for training (4688793 samples) and 10% for both validation and testing (with 532340 and 537141 samples, respectively). As for the finetuning dataset, we isolated a total of 1.5M training samples and 0.17M samples, both for validation and test.

For both training tasks, the parameters were optimized using the Adam optimizer [47] with *β*_1_ = 0.9, *β*_2_ = 0.999 and *ϵ* = 10^−9^. The learning rate was varied over the course of training, following the same schedule described in [42], with the following parameters: warmup_steps of 2000 and 3000, for base and large models respectively, and a factor of 0.1, during pre-training; during fine-tuning, warmup_steps and factor were set to 2000 and 0.01, respectively, for both model configurations. Additionally, we applied gradient clipping to the model’s parameters, limiting the maximum L2 norm of the gradients to 1, which helps prevent exploding gradients and promotes a more stable training of the neural network. The batch size was set to 5, due to computational and memory restrictions. However, since the model makes predictions on the token level and different batches can vary in terms of sequence length, the true batch size used for loss computation is actually higher. As the loss function, we used cross-entropy (CE) loss for all tasks and model configurations, except for the fine-tuning of the base model. In the latter, a binary cross-entropy (BCE) loss was implemented for each one of the three target classes. Furthermore, as the pre-training dataset was imbalanced, we set class weights inversely proportional to the respective nucleotide frequency in the data. Dropout Layers (with p=0.1) were added, one after the Tokenizer and Embedder’s output, another before the final Linear Layer of every task head and also in the transformer encoder, as described in [42]. These hyperparameters were also established empirically and based on the literature.

We pre-trained and fine-tuned the base model for 30 epochs and the large one for 5 and 9, respectively. Metrics such as accuracy, precision, recall and f1-score were used to evaluate the models’ performances.

#### Training hardware

All experiments with the base model configuration were run in a NVIDIA RTX 4090 GPU (with 24GB GDDR6X memory). For the large model, we used a cloud-based GPU service, specifically RunPod, utilizing an NVIDIA A100 GPU with 80GB of GDDR6X memory. Models were developed with python 3.11.4 and Pytorch 2. We used the Automatic Mixed Precision package from pytorch to implement mixed precision training, enabling faster computation and reduced memory usage on GPU while maintaining training stability and accuracy.

### Self-Attention analyses

Self-attention in transformers allows the model to relate different parts of a sequence by generating queries, keys, and values for each position. The model compares each query with all keys in the sequence to find relevant information based on context, effectively retrieving information from within the sequence itself. This dynamic process helps the model focus on important patterns and relationships between elements, and is captured in attention weight matrices that show how much attention each position in the sequence pays to others.

We evaluated these attention weight matrices to validate our model’s performance by examining the patterns and dependencies it captures. Our goal was to ensure that C.La.P accurately leverages biological context and key regions of the genomic sequence in its predictions. This analysis was conducted on the fine-tuned model’s best epoch (the one with lower validation loss), as it is specifically designed to predict relevant epigenetic marks in genomic sequences. Our first objective was to determine where the model focused the most when predicting a CTCF region in the input sample. To achieve this, we applied the attention-rollout technique [33], which aggregates the attention matrices throughout all the layers and heads of the encoder, producing a sample-wide array of attention values. These values were then normalized for each sample, ranging from 0 and 1. To detect positions where the base model pays the most attention, we simply choose the position with the highest value. About 95% of these mA positions overlap with CTCF PWM hits (See Methods-PWMs). Doing the same analysis with the large model results in a much lower overlap with CTCF PWM hits, even though empirically we can still see the model’s attention peaking over those. To refine the large model’s attention rollout signal, we applied a moving average filter with a window size of 5, smoothing out isolated noise and highlighting more robust regions of significant attention. We then used a method called peak-calling based on scipy’s find_peaks function (scipy.signal.find_peaks in scipy version 1.10.1). After smoothing and peak-calling, the large model’s mA positions show an overlap of about 85% with CTCF PWM hits showing that indeed the large model’s attention also tends to peak over those genomic features.

### PWMs

Positional Weight Matrices (PWMs) [48], are mathematical models used to represent TFbs and quantify the sequence preferences of these DNA-binding proteins, like CTCF. A higher PWM score indicates a greater likelihood that the nucleotide sequence in that region corresponds to a TFbs. We used genome wide CTCF hits as computed in [49].

