## Supplementary figures and images for "C.La.P.: Enhancing transformer-based genomic signal modeling by integrating DNA sequences and chromatin accessibility data"

### Supplemental CTCF examples

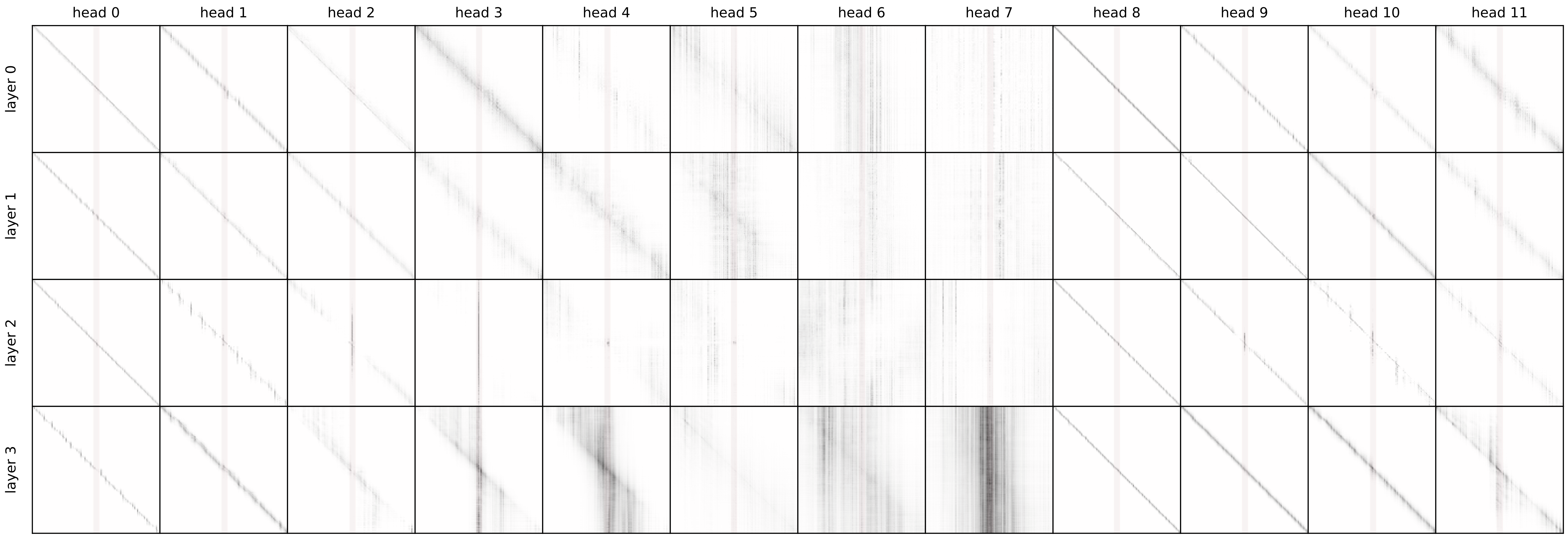

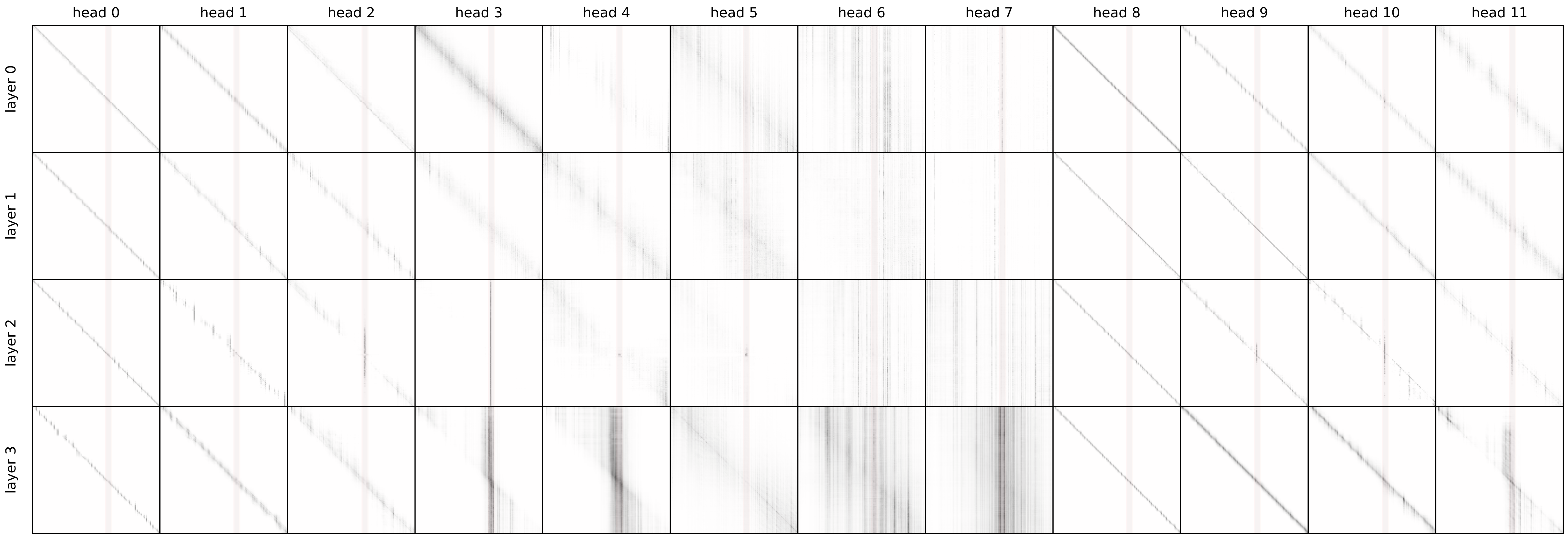

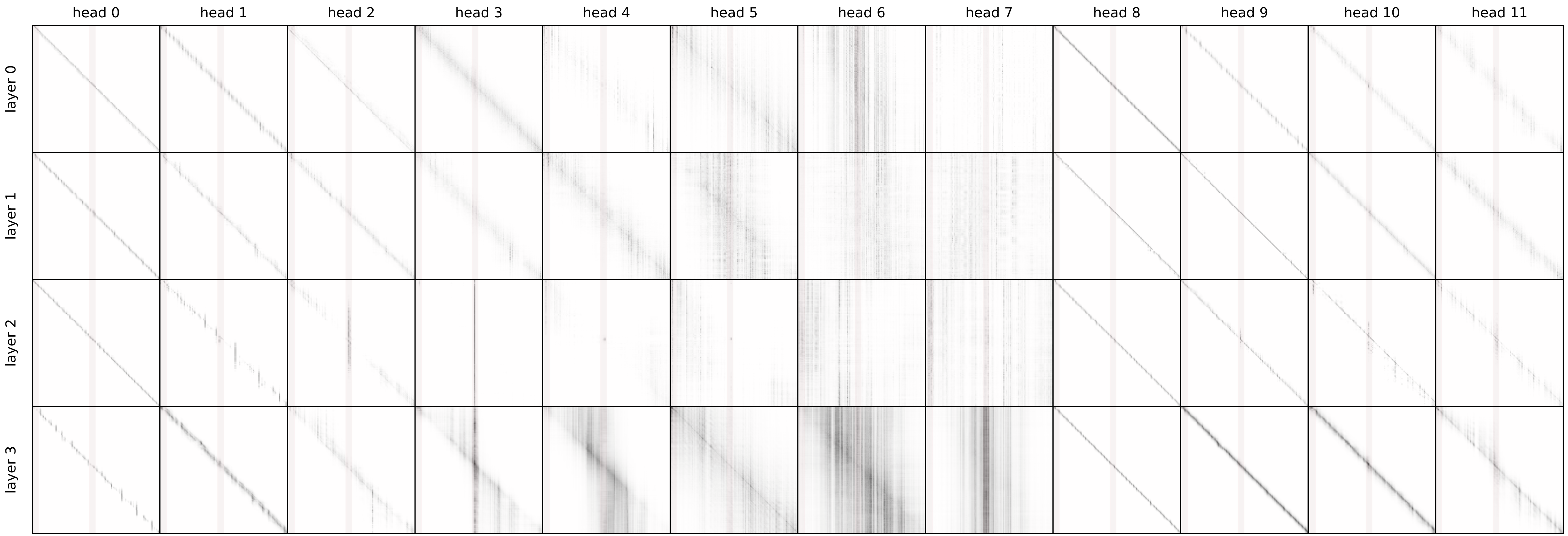

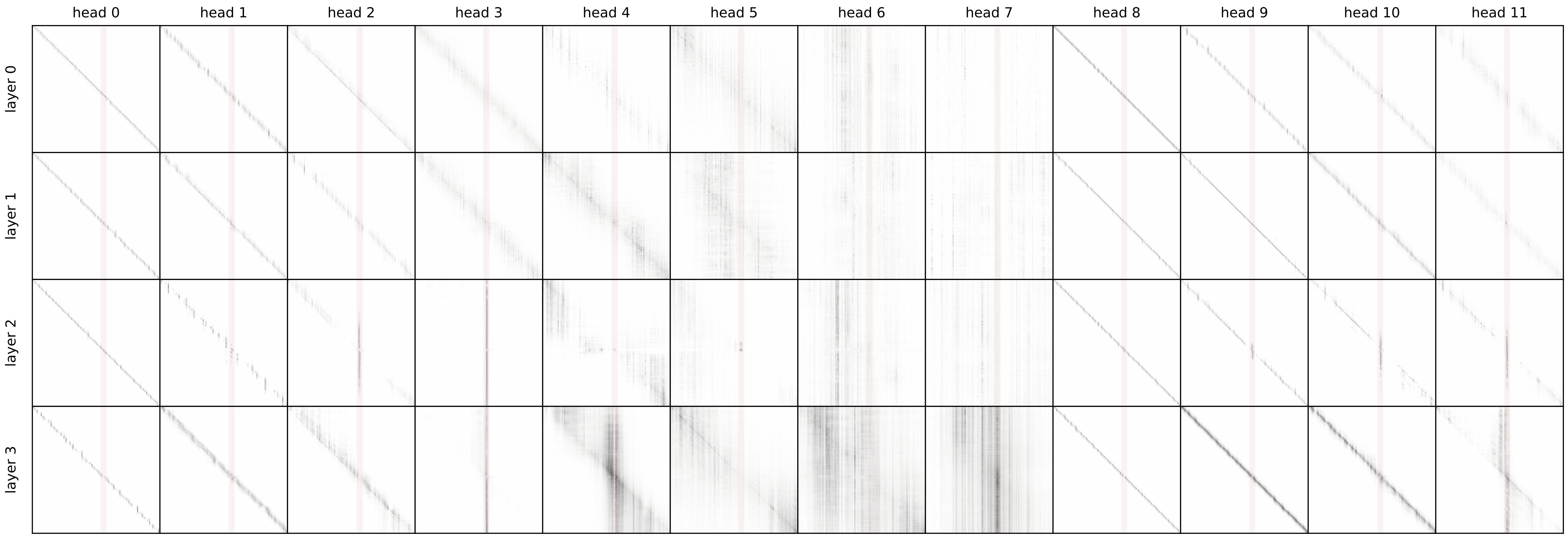

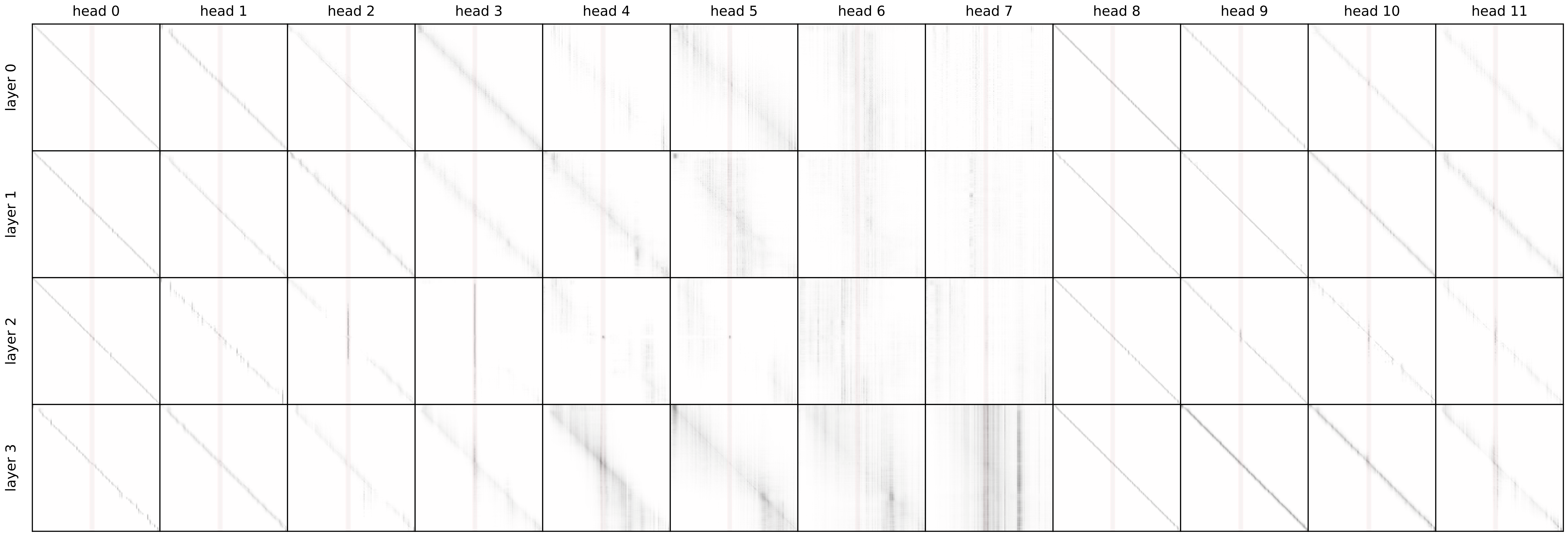

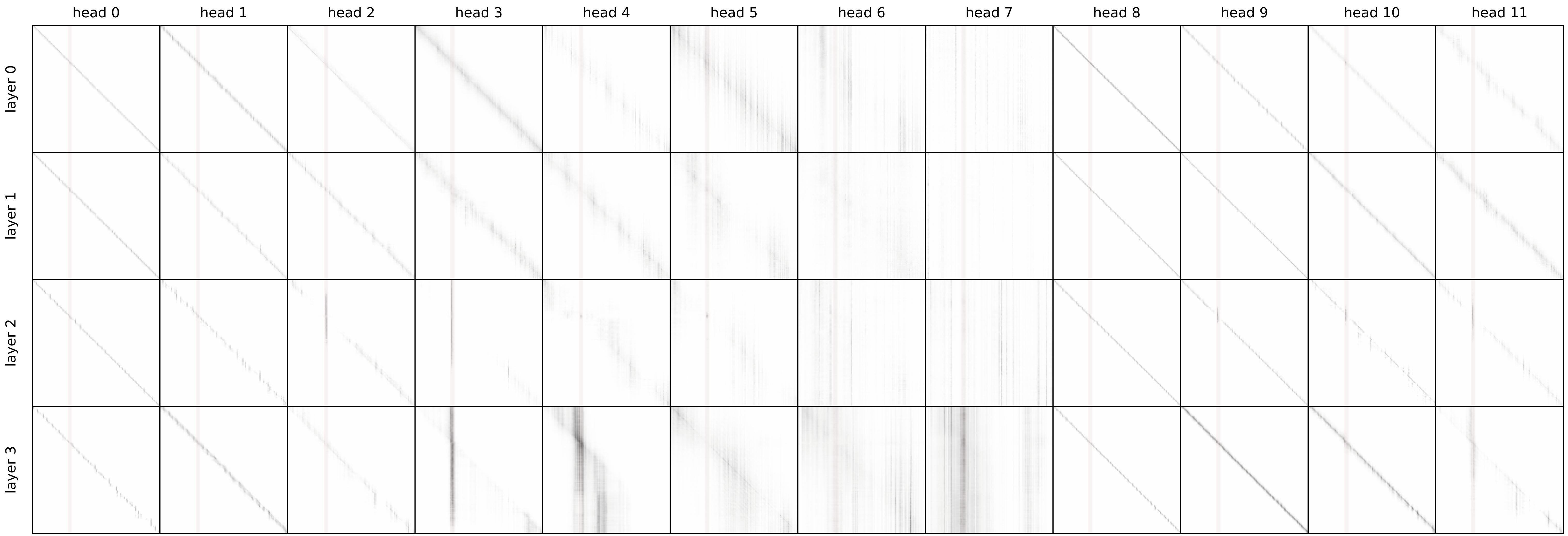

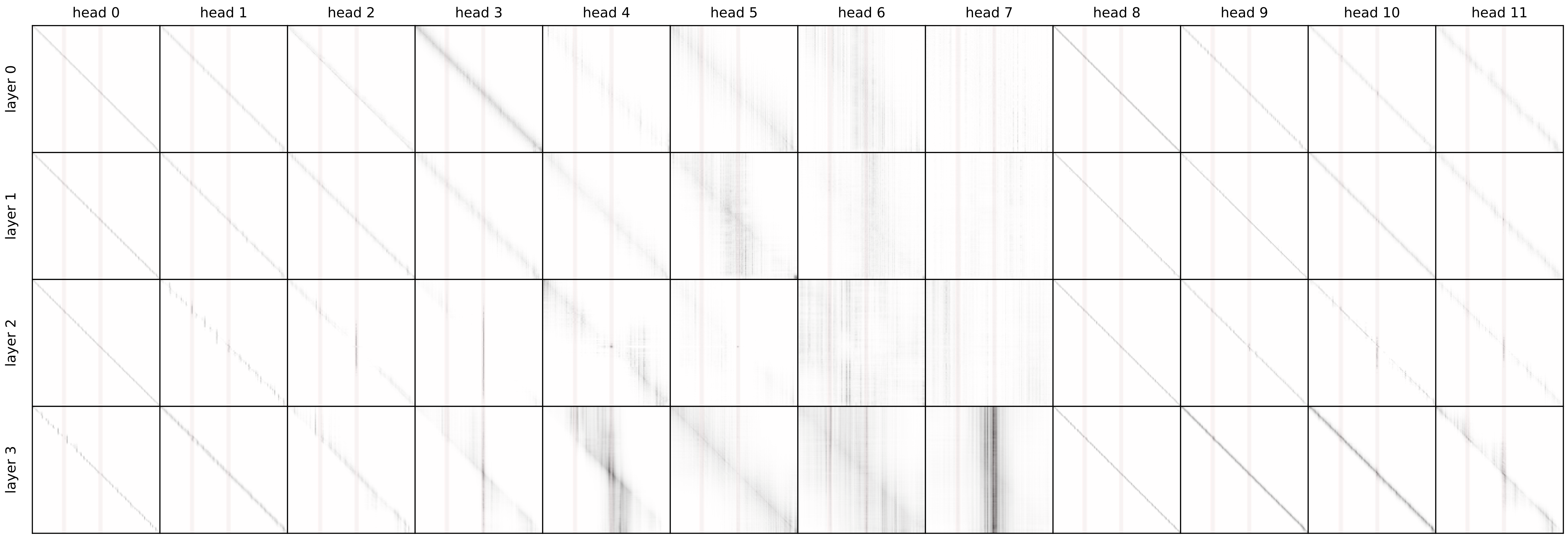

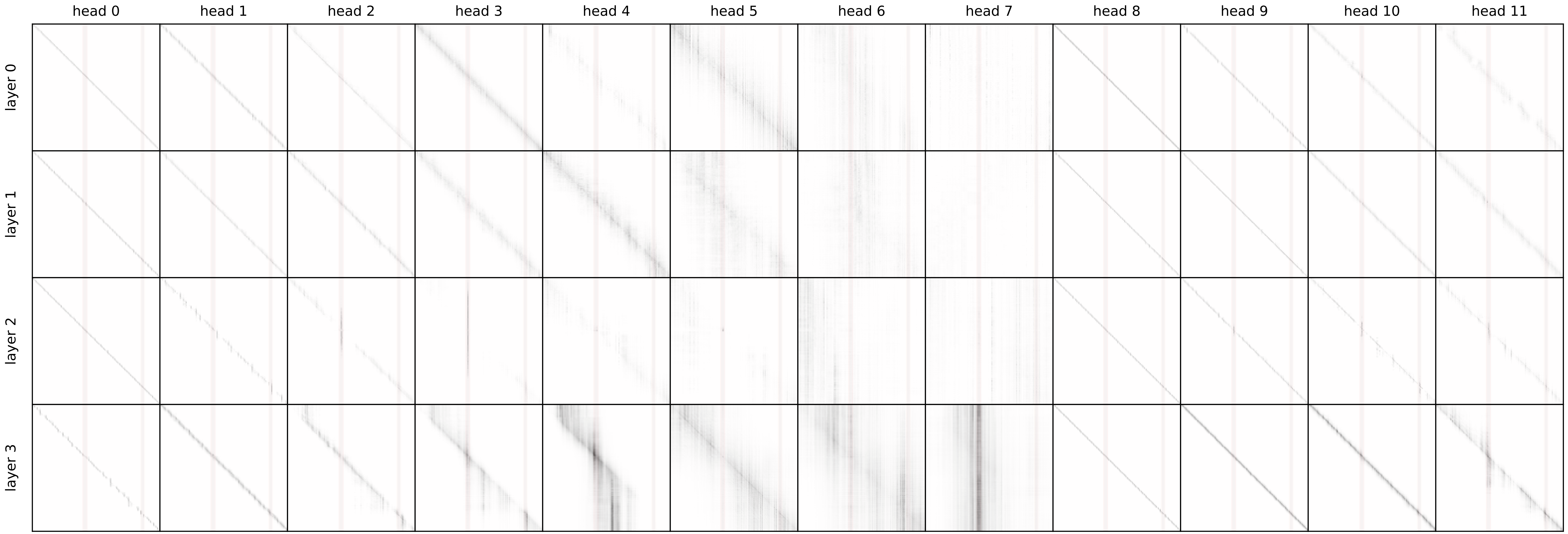

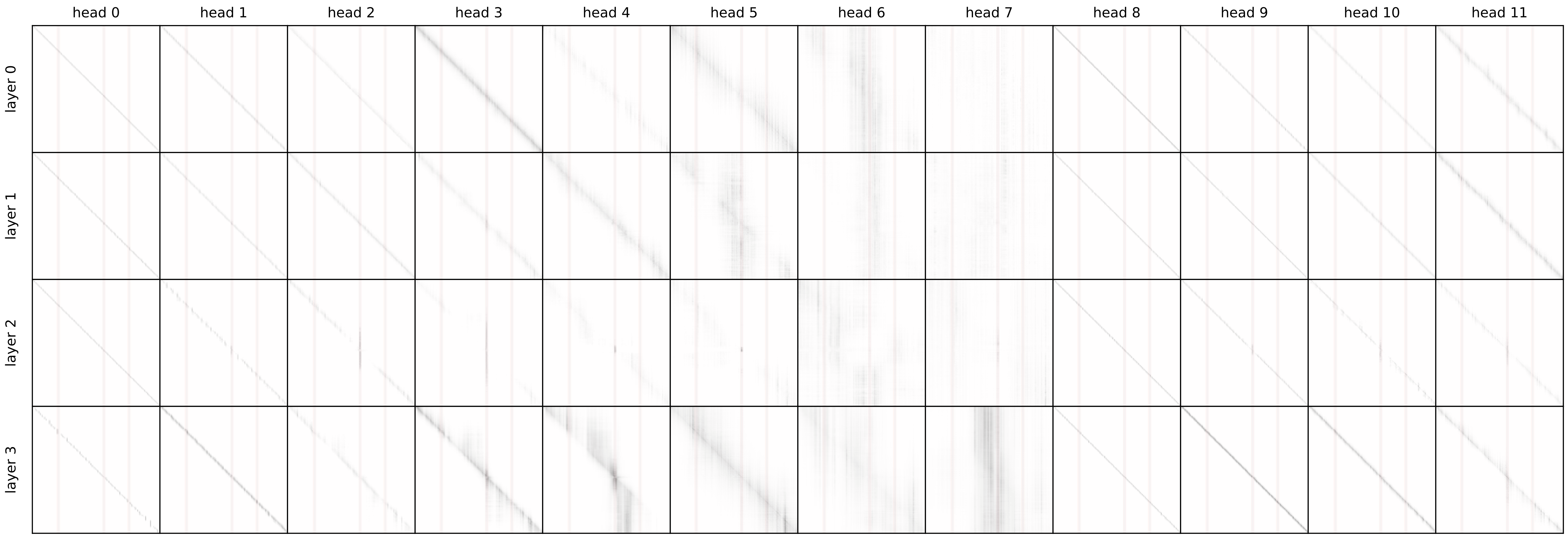

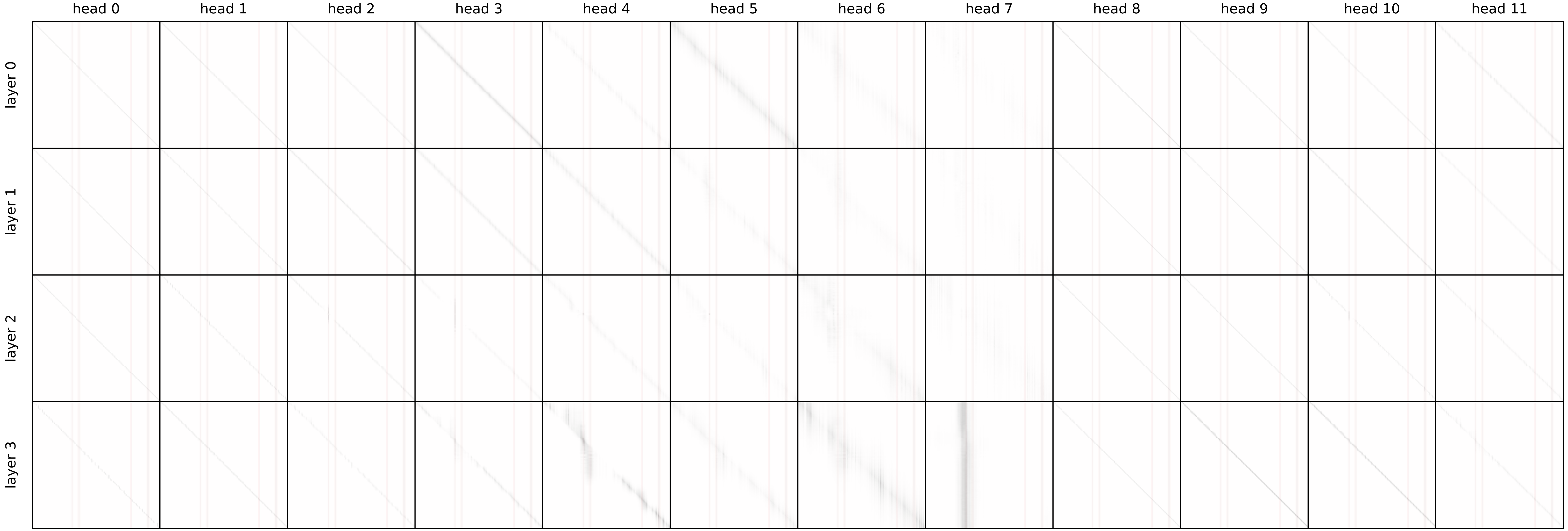
