## Supplemental Figures for "C.La.P.: Enhancing transformer-based genomic signal modeling by integrating DNA sequences and chromatin accessibility data"

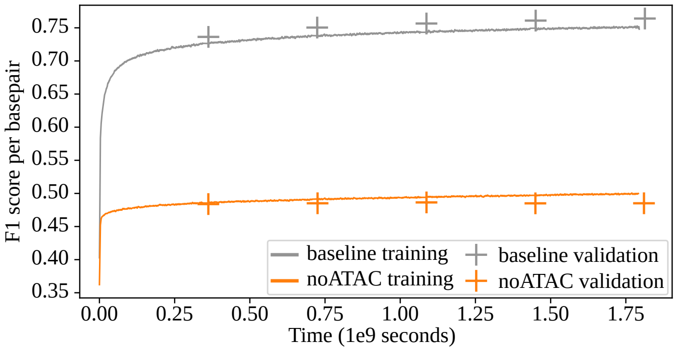


**Figure 1:** ***Pre-training without ATAC-seq signal*** *In the experiment plotted with a yellow line here, we attempted to pre-train our model while withholding the two ATAC-seq features of the input array. The model fails to 'take off' in these conditions, showcasing the importance of the ATAC-seq signal. Without these features the model is essentially a sequence-only model.*

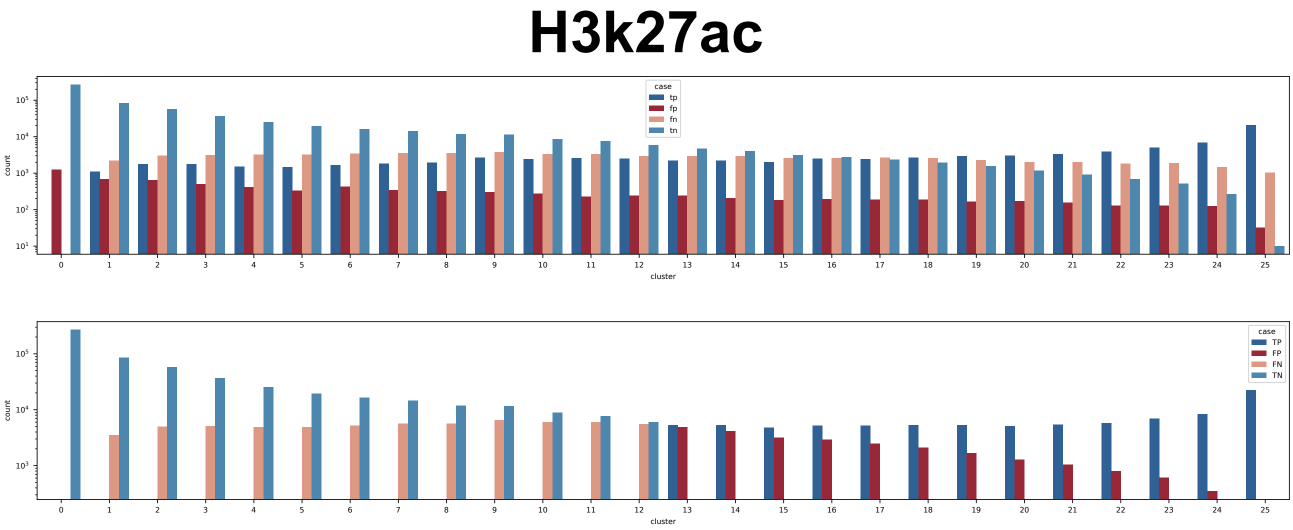


**Figure 2:** *Clusters from the bd-clustered dataset were grouped on the x-axis by the number of tissues (0–25) in which they are active for H3K27ac. The y-axis (log scale) shows TP, FP, FN, and TN predictions per group for CLaP and a hypothetical sequence-only classifier, as detailed in the main text.*

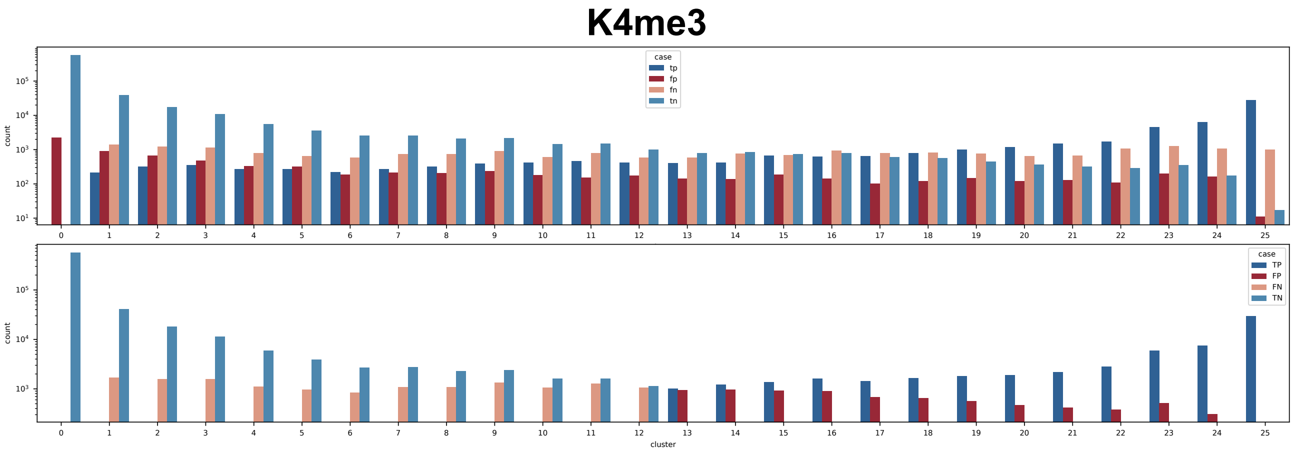


**Figure 3:** *Clusters from the bd-clustered dataset were grouped on the x-axis by the number of tissues (0–25) in which they are active for H3K4me3. The y-axis (log scale) shows TP, FP, FN, and TN predictions per group for CLaP and a hypothetical sequence-only classifier, as detailed in the main text.*

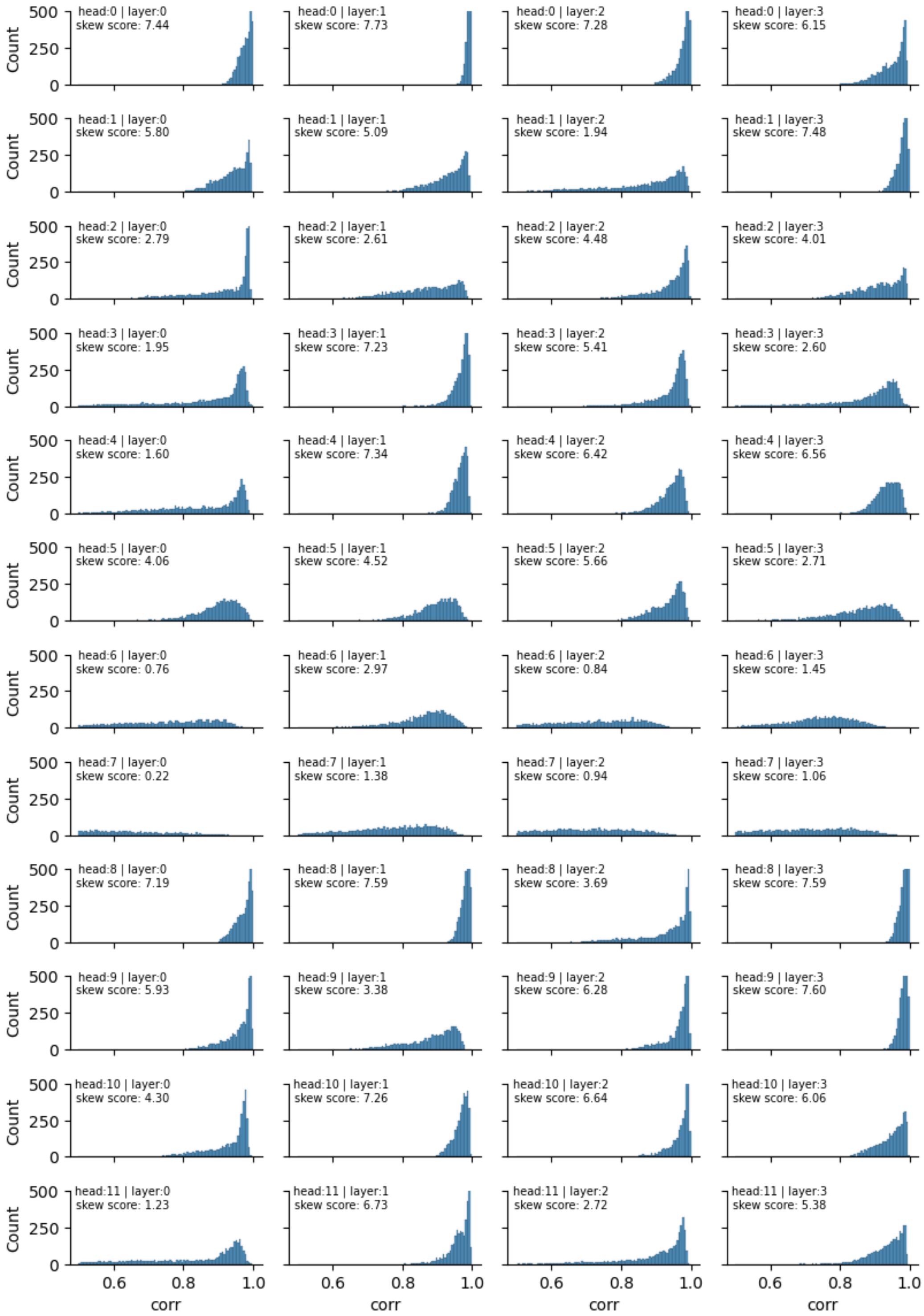


**Figure 4:** *We computed the correlations of each head's self-attention matrix, between pairs of samples from different biosamples, where one sample was poor in ATAC-seq signal and the other sample was rich. These samples have the exact same reference sequence, so high correlations signify that the head is producing the same output between the two samples and thus its activation is very reliant on the genomic sequence. Low correlation values, on the other hand, signify that the head reacts differently between the two samples, thus its activation is more reliant on the values of the ATAC-seq signal.*

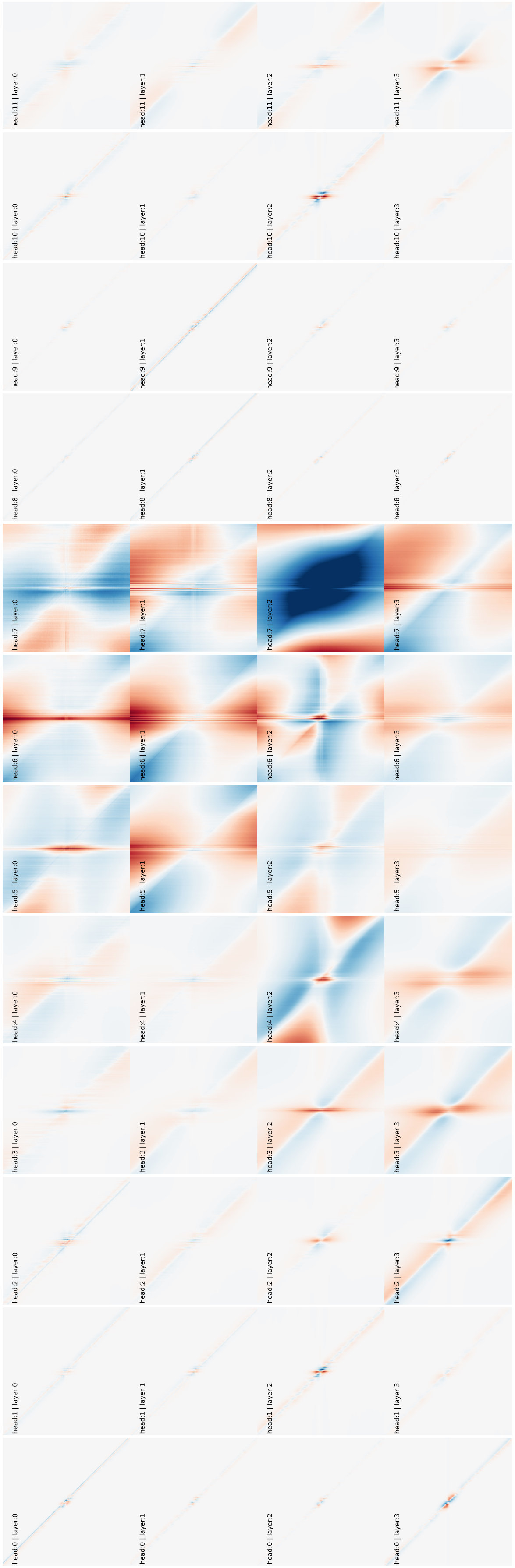


**Figure 5:** *We collected a number of CTCF PWM hits that fall inside CTCF ChIPseq ground truth regions. We then register the values of each head of the model for this collection of genomic positions, and normalize by subtracting the values of each head for an equally big collection of random genomic positions. This way we can visualize how each head reacts when it encounters a CTCF site versus background noise.*

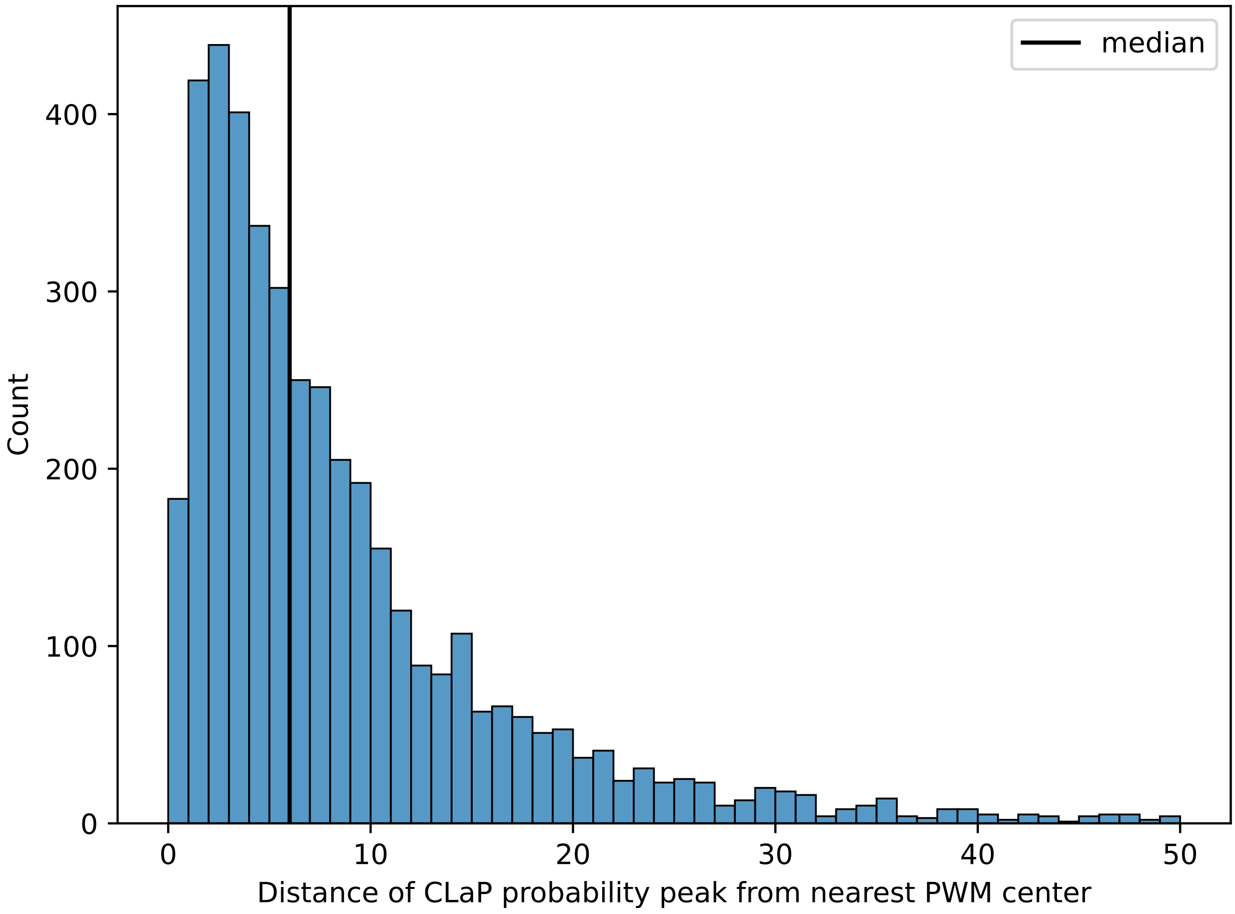


**Figure 6:** *A histogram of the distances of CLP peaks from the nearest PWM hit center.*

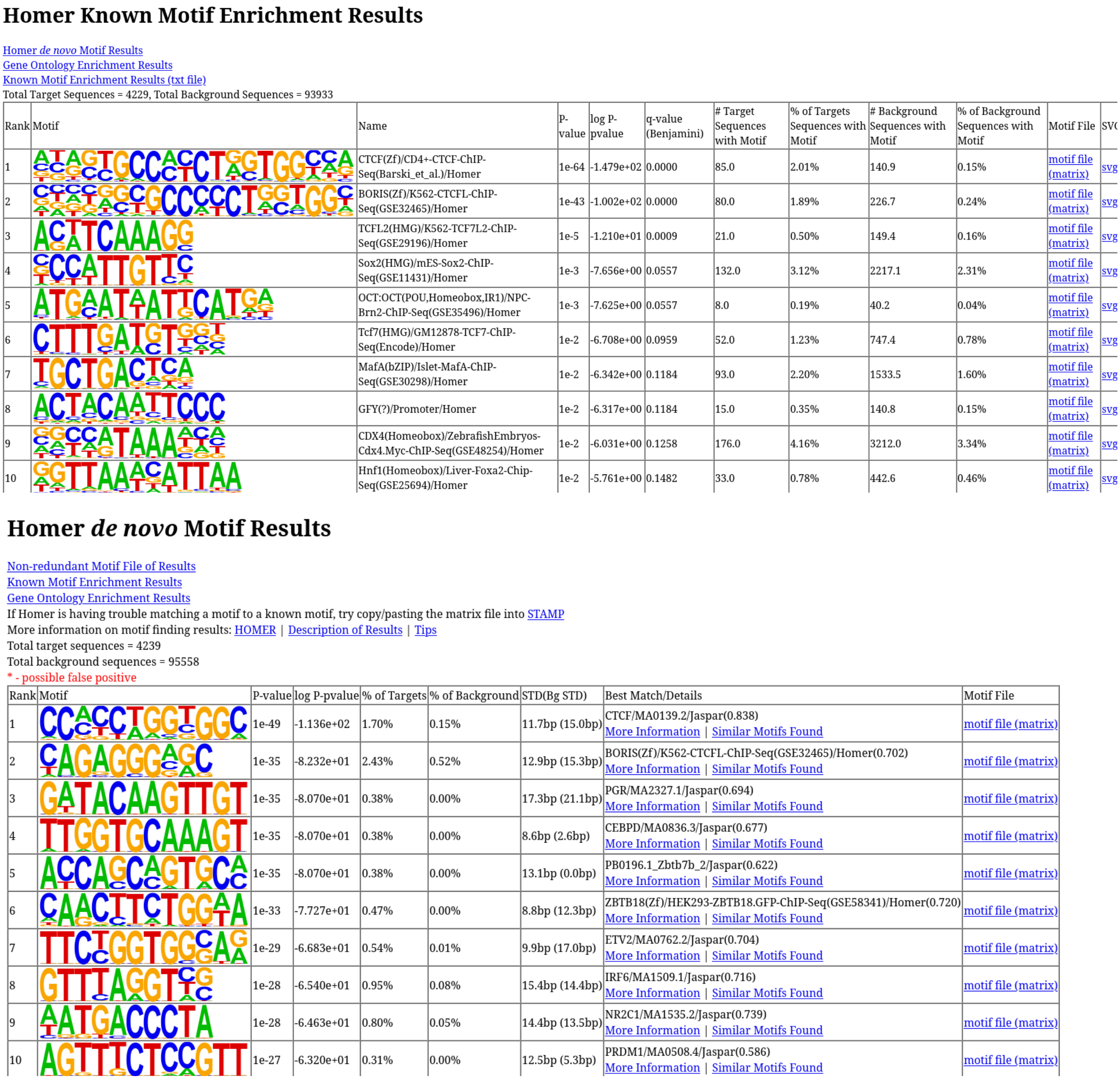

### Homer Known Motif Enrichment Results

Homer *de novo* Motif Results
Gene Ontology Enrichment Results
Known Motif Enrichment Results (txt file)
Total Target Sequences = 4229, Total Background Sequences = 93933

| Rank | Motif | Name | P-value | log P-pvalue | q-value (Benjamini) | # Target Sequences with Motif | % of Targets Sequences with Motif | # Background Sequences with Motif | % of Background Sequences with Motif | Motif File | SV |
|---|---|---|---|---|---|---|---|---|---|---|---|
| 1 | | CTCF(Zf)/CD4+-CTCF-ChIP-Seq(Barski_et_al.)/Homer | 1e-64 | -1.479e+02 | 0.0000 | 85.0 | 2.01% | 140.9 | 0.15% | motif file (matrix) | svg |
| 2 | | BORIS(Zf)/K562-CTCFL-ChIP-Seq(GSE32465)/Homer | 1e-43 | -1.002e+02 | 0.0000 | 80.0 | 1.89% | 226.7 | 0.24% | motif file (matrix) | svg |
| 3 | | TCFL2(HMG)/K562-TCF7L2-ChIP-Seq(GSE29196)/Homer | 1e-5 | -1.210e+01 | 0.0009 | 21.0 | 0.50% | 149.4 | 0.16% | motif file (matrix) | svg |
| 4 | | Sox2(HMG)/mES-Sox2-ChIP-Seq(GSE11431)/Homer | 1e-3 | -7.656e+00 | 0.0557 | 132.0 | 3.12% | 2217.1 | 2.31% | motif file (matrix) | svg |
| 5 | | OCT:OCT(POU,Homeobox,IR1)/NPC-Brn2-ChIP-Seq(GSE35496)/Homer | 1e-3 | -7.625e+00 | 0.0557 | 8.0 | 0.19% | 40.2 | 0.04% | motif file (matrix) | svg |
| 6 | | Tcf7(HMG)/GM12878-TCF7-ChIP-Seq(Encode)/Homer | 1e-2 | -6.708e+00 | 0.0959 | 52.0 | 1.23% | 747.4 | 0.78% | motif file (matrix) | svg |
| 7 | | MafA(bZIP)/Islet-MafA-ChIP-Seq(GSE30298)/Homer | 1e-2 | -6.342e+00 | 0.1184 | 93.0 | 2.20% | 1533.5 | 1.60% | motif file (matrix) | svg |
| 8 | | GFY(?)/Promoter/Homer | 1e-2 | -6.317e+00 | 0.1184 | 15.0 | 0.35% | 140.8 | 0.15% | motif file (matrix) | svg |
| 9 | | CDX4(Homeobox)/ZebrafishEmbryos-Cdx4.Myc-ChIP-Seq(GSE48254)/Homer | 1e-2 | -6.031e+00 | 0.1258 | 176.0 | 4.16% | 3212.0 | 3.34% | motif file (matrix) | svg |
| 10 | | Hnf1(Homeobox)/Liver-Foxa2-Chip-Seq(GSE25694)/Homer | 1e-2 | -5.761e+00 | 0.1482 | 33.0 | 0.78% | 442.6 | 0.46% | motif file (matrix) | svg |

### Homer *de novo* Motif Results

Non-redundant Motif File of Results
Known Motif Enrichment Results
Gene Ontology Enrichment Results
If Homer is having trouble matching a motif to a known motif, try copy/pasting the matrix file into STAMP
More information on motif finding results: HOMER | Description of Results | Tips
Total target sequences = 4239
Total background sequences = 95558
* - possible false positive

| Rank | Motif | P-value | log P-pvalue | % of Targets | % of Background | STD(Bg STD) | Best Match/Details | Motif File |
|---|---|---|---|---|---|---|---|---|
| 1 | | 1e-49 | -1.136e+02 | 1.70% | 0.15% | 11.7bp (15.0bp) | CTCF/MA0139.2/Jaspar(0.838) More Information \| Similar Motifs Found | motif file (matrix) |
| 2 | | 1e-35 | -8.232e+01 | 2.43% | 0.52% | 12.9bp (15.3bp) | BORIS(Zf)/K562-CTCFL-ChIP-Seq(GSE32465)/Homer(0.702) More Information \| Similar Motifs Found | motif file (matrix) |
| 3 | | 1e-35 | -8.070e+01 | 0.38% | 0.00% | 17.3bp (21.1bp) | PGR/MA2327.1/Jaspar(0.694) More Information \| Similar Motifs Found | motif file (matrix) |
| 4 | | 1e-35 | -8.070e+01 | 0.38% | 0.00% | 8.6bp (2.6bp) | CEBPD/MA0836.3/Jaspar(0.677) More Information \| Similar Motifs Found | motif file (matrix) |
| 5 | | 1e-35 | -8.070e+01 | 0.38% | 0.00% | 13.1bp (0.0bp) | PB0196.1_Zbtb7b_2/Jaspar(0.622) More Information \| Similar Motifs Found | motif file (matrix) |
| 6 | | 1e-33 | -7.727e+01 | 0.47% | 0.00% | 8.8bp (12.3bp) | ZBTB18(Zf)/HEK293-ZBTB18.GFP-ChIP-Seq(GSE58341)/Homer(0.720) More Information \| Similar Motifs Found | motif file (matrix) |
| 7 | | 1e-29 | -6.683e+01 | 0.54% | 0.01% | 9.9bp (17.0bp) | ETV2/MA0762.2/Jaspar(0.704) More Information \| Similar Motifs Found | motif file (matrix) |
| 8 | | 1e-28 | -6.540e+01 | 0.95% | 0.08% | 15.4bp (14.4bp) | IRF6/MA1509.1/Jaspar(0.716) More Information \| Similar Motifs Found | motif file (matrix) |
| 9 | | 1e-28 | -6.463e+01 | 0.80% | 0.05% | 14.4bp (13.5bp) | NR2C1/MA1535.2/Jaspar(0.739) More Information \| Similar Motifs Found | motif file (matrix) |
| 10 | | 1e-27 | -6.320e+01 | 0.31% | 0.00% | 12.5bp (5.3bp) | PRDM1/MA0508.4/Jaspar(0.586) More Information \| Similar Motifs Found | motif file (matrix) |

**Figure 7:** *Homer analysis in regions of 60bps around CLP peaks. Both de-novo and known motifs show a clear enrichment for the CTCF PWMs*

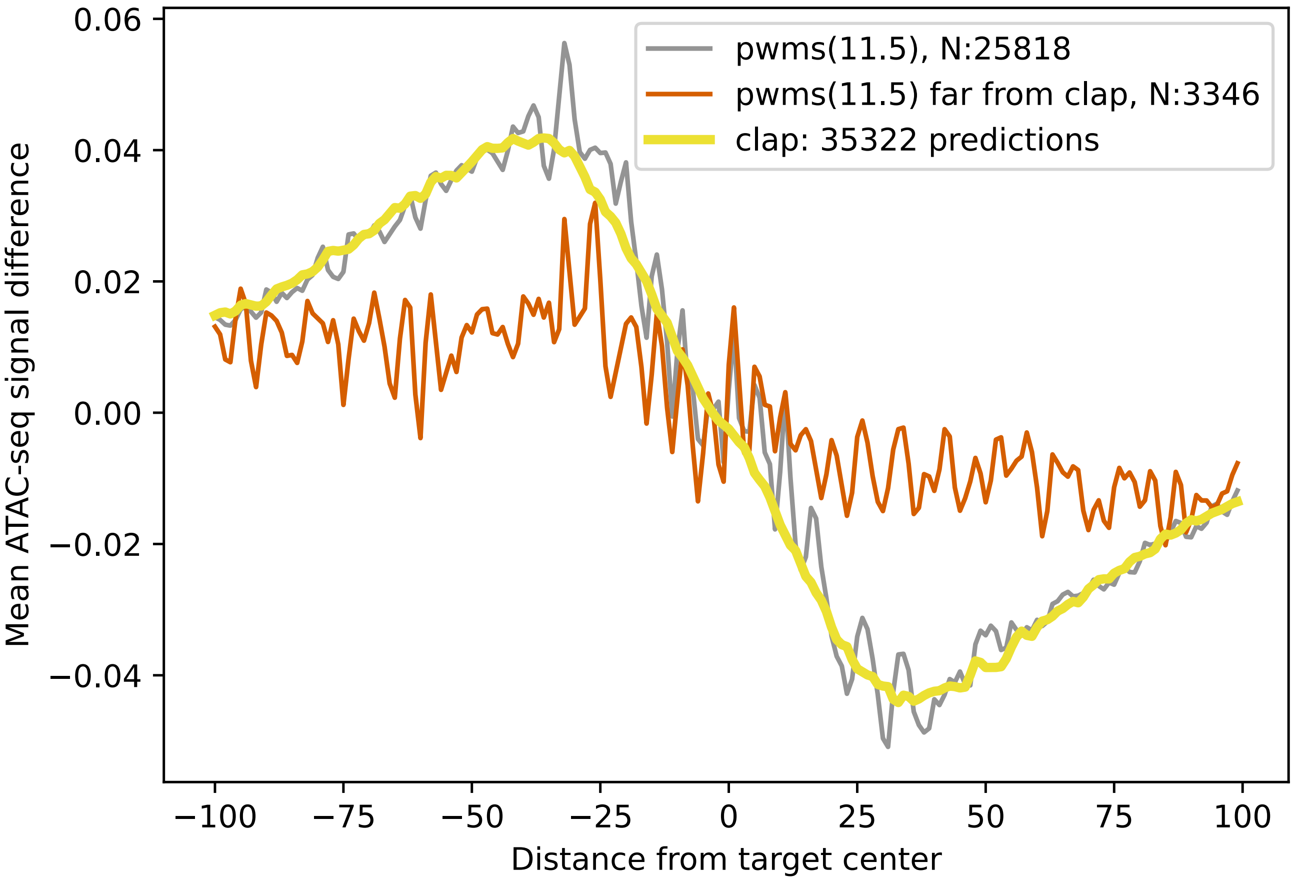


**Figure 8:** *Plotting the normalized difference between "+" and "-" ATAC-seq signals allows us to compare sets of positions. Here we compare CLP peaks and PWM hits with score higher than 11.5, both found inside the samples of our validation set and overlapping with ground truth regions, making them 'True Positives'. The third set, PWMs far from CLAP, is a subset of the PWM set containing only the PWMs that did not overlap a CLP peak. The CLaP set maintains comparable Atac-seq imbalance (ASI) to the PWM set, while having about 36% more sites. Furthermore, the subset of PWMs that does not overlap CLP positions has a much weaker ASI, suggesting that many of these predictions are not real binding sites.*
